# Mechanism of ribosome-associated mRNA degradation during tubulin autoregulation

**DOI:** 10.1101/2023.01.12.523749

**Authors:** Markus Höpfler, Eva Absmeier, Sew-Yeu Peak-Chew, Evangelia Vartholomaiou, Lori A. Passmore, Ivana Gasic, Ramanujan S. Hegde

**Affiliations:** MRC Laboratory of Molecular Biology, Cambridge, CB2 0QH, United Kingdom; Department of Cell Biology, University of Geneva, Geneva, Switzerland

## Abstract

**SUMMARY:** Microtubules play crucial roles in cellular architecture, intracellular transport, and mitosis. The availability of free tubulin subunits impacts polymerization dynamics and microtubule function. When cells sense excess free tubulin, they trigger degradation of the encoding mRNAs, which requires recognition of the nascent polypeptide by the tubulin-specific ribosome-binding factor TTC5. How TTC5 initiates decay of tubulin mRNAs is unknown. Here, our biochemical and structural analysis reveals that TTC5 recruits the poorly studied protein SCAPER to the ribosome. SCAPER in turn engages the CCR4-NOT deadenylase complex through its CNOT11 subunit to trigger tubulin mRNA decay. SCAPER mutants that cause intellectual disability and retinitis pigmentosa in humans are impaired in CCR4-NOT recruitment, tubulin mRNA degradation, and microtubule-dependent chromosome segregation. Our findings demonstrate how recognition of a nascent polypeptide on the ribosome is physically linked to mRNA decay factors via a relay of protein-protein interactions, providing a paradigm for specificity in cytoplasmic gene regulation.

## INTRODUCTION

Microtubules (MTs) constitute a fundamental part of the eukaryotic cytoskeleton with key roles in shaping widely varying cellular architectures, in facilitating transport within cells over long distances, and in segregating chromosomes during cell division^1,2^. These functions rely on the highly dynamic assembly and disassembly of MTs from heterodimeric subunits comprising a- and β-tubulins^1,2^. Microtubules are regulated by more than 40 microtubule-associated proteins (MAPs) that modify the behaviour of individual MTs and their assembly into higher-order structures^3^. Furthermore, tubulins are subject to an extensive range of post-translational modifications, some of them exclusively found on tubulins^4,5^.

Despite research on tubulin and MTs for many decades, several crucial MT regulators have only recently been identified and are often still poorly characterized^6–8^. Many of these are linked to human pathologies such as cancer, neurodevelopmental or neurodegenerative conditions, and represent potential targets for therapeutics that could complement other tubulin-targeting drugs like taxol and colchicine^8–12^. Thus, accurate MT regulation is of exceptionally broad importance and deciphering the range of pathways that impinge on tubulins is crucial for understanding and modulating the progression of various pathologic states.

A crucial parameter for the balance between MT growth and shrinkage is the concentration of the free tubulin subunits^1,13^. Several decades ago, it was recognized that cellular tubulin concentration is tightly controlled in part by a feedback mechanism termed tubulin autoregulation^14–16^. This widely conserved phenomenon dynamically adjusts tubulin mRNA levels in response to changes in the level of free tubulin subunits. Regulation occurs strictly post-transcriptionally and involves translation-dependent mRNA degradation that is preferentially triggered under conditions of excess free tubulin. How this highly selective autoregulatory loop operates has long been mysterious.

The only known component in the tubulin autoregulation pathway is TTC5, a recently discovered factor that recognizes an N-terminal peptide motif common to nascent tubulin polypeptides emerging from translating ribosomes^17^. Mutations that impair TTC5 recognition of tubulin ribosome-nascent-chains (RNCs) abolish autoregulation and lead to aberrant mitosis^17^, a highly sensitive measure of perturbed microtubule dynamics^18,19^. Although the discovery and validation of TTC5 finally provided a molecular handle for the tubulin autoregulation pathway, it is not known why TTC5 binding at the polypeptide exit tunnel of tubulin-producing ribosomes leads to degradation of the associated mRNAs. Furthermore, the broader biological relevance of autoregulation for human physiology is unclear.

While multiple cases of mRNA sequence-dependent post-transcriptional regulation have been well characterized^20–22^, the molecular basis for the coupling of nascent chain recognition to selective mRNA degradation is poorly understood. Prominent examples of such nascent peptide-dependent regulation include highly expressed mRNAs such as those coding for endoplasmic reticulum-targeted proteins^23,24^ and ribosomal proteins^25^. Recent studies suggest that bacteriophage-derived anti-CRISPR proteins recognize nascent Cas12a protein to trigger degradation of Cas12a mRNA, suggesting related mechanisms exist beyond eukaryotes^26,27^.

Conceptually, specific nascent peptide recognition coupled to mRNA decay is reminiscent of the well-studied co-translational capture of signal sequences by the signal recognition particle that elicits RNC targeting to the endoplasmic reticulum (ER) membrane. Similarly, nascent chains can be used to direct mRNAs to other locations such as centrosomes, apical poles in epithelial cells during embryogenesis and others^28–31^, or to drive mRNA co-localization for co-translational protein complex assembly^32–33^. These examples illustrate that nascent chain-directed mRNA fate decisions are broadly relevant, but the molecular mechanisms and structural features linking peptide recognition to downstream events are enigmatic in most cases. Given the critical functions of microtubules in numerous areas of cell and organism homeostasis^18,34^, neuronal cell function^12^, and their relevance as drug targets^9,10^, we sought to understand the mechanistic basis of co-translational mRNA decay using tubulins as an example.

## RESULTS

### TTC5 recruits SCAPER to ribosomes

Tubulin autoregulation can be experimentally induced by microtubule depolymerizing drugs such as colchicine or combretastatin A4 (CA4). The acute rise in free tubulin heterodimers comprising α and β subunits liberates TTC5 from a yet-unidentified sequestration factor^17^. TTC5 then engages tubulin-synthesizing ribosomes and triggers degradation of tubulin mRNAs by an unknown process. To identify factors downstream of ribosome-bound TTC5, we used a biotin proximity labelling strategy^35^. The promiscuous biotin ligase TurboID was fused to TTC5 to biotinylate interaction partners during ongoing tubulin mRNA degradation (Fig. 1A). As a specificity control, we identified a mutation within a conserved surface residue of TTC5 (K97A; Fig. S1A) that abolished TTC5’s capacity to trigger tubulin mRNA degradation (Fig. 1B; Fig. S1B) despite normal expression (Fig. S1C) and unimpaired recruitment to tubulin RNCs (Fig. S1D).

**Figure 1.**
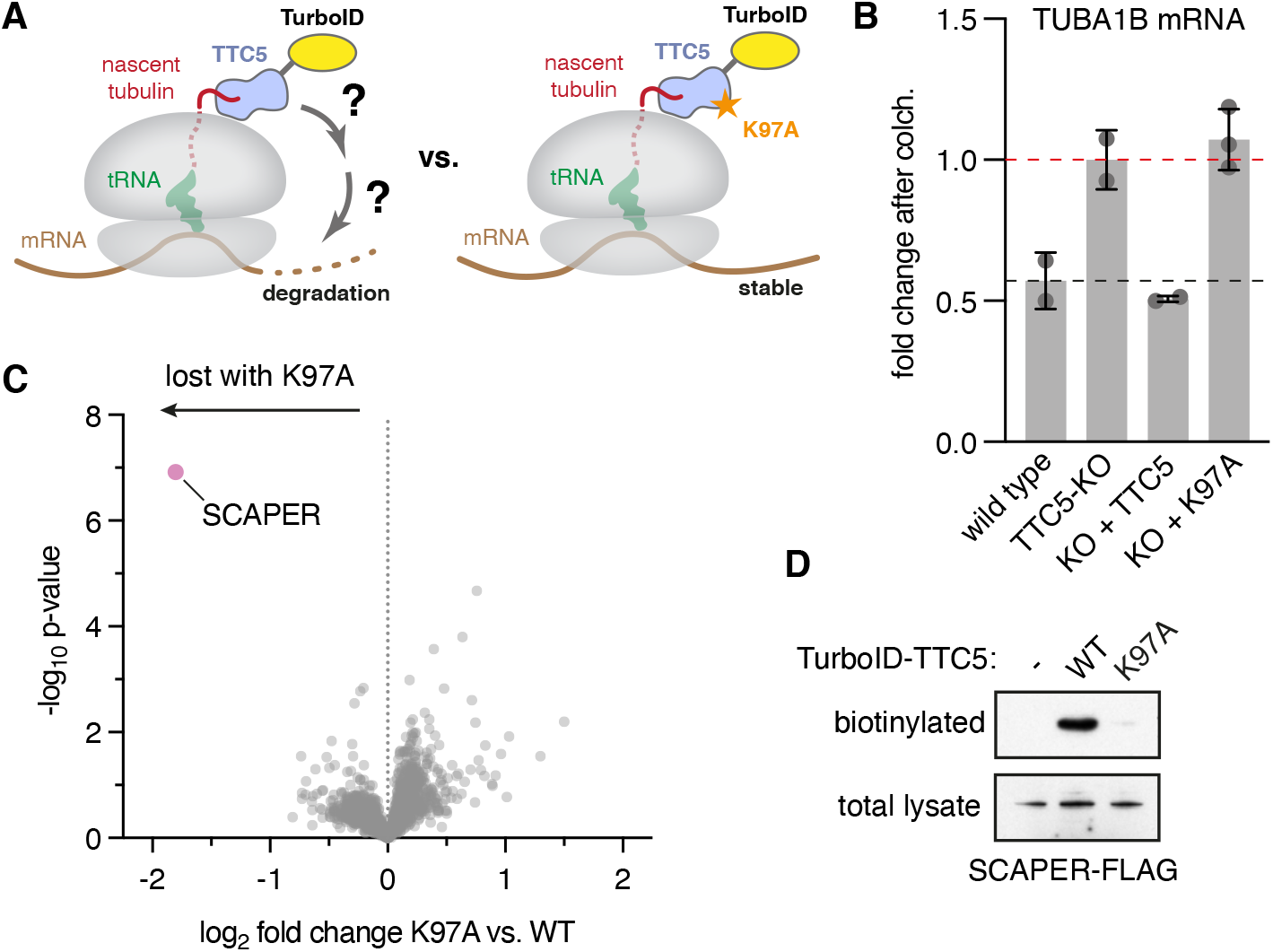
TTC5 proximity labelling identifies SCAPER as autoregulation-dependent interactor. (A) Strategy for identification of tubulin autoregulation factors acting downstream of TTC5. Proximity labelling was achieved by fusing TurboID to the N-terminus of either wild type TTC5 or the Lys^97^→Ala (K97A) mutant. (B) Quantification of tubulin mRNA in HEK293 T-REx cells by reverse transcription followed by quantitative real-time PCR. The fold change of TUBA1B mRNA after 3h 10 μM colchicine (colch.) treatment is plotted (mean +/− standard deviation from 2 replicates, 3 for K97A). This is hereafter referred to as the “autoregulation assay”. The red dashed line indicates the starting tubulin mRNA level prior to colchicine, arbitrarily set to a value of 1. The black dashed line indicates the fold change in wild type (WT) cells. This is typically ~0.5 after 3h of colchicine, reflective of 50% mRNA degradation, but varies slightly in different experiments due to minor variations in experimental conditions. TTC5 knockout (KO) was complemented by re-expressing GFP-tagged wild type or K97A TTC5. (C) Proximity labelling using TurboID fused to either wild type (WT) or mutant (K97A) TTC5 followed by enrichment of biotinylated proteins and quantitative mass spectrometry. See also Supplementary Table S2. (D) Proximity labelling assay as in panel C with overexpression of FLAG-tagged SCAPER in the indicated cell lines. Total lysates were probed with anti-FLAG antibody and the biotinylated population with anti-SCAPER antibody. Endogenous SCAPER is not detected at this exposure due to its low expression.

After confirming that the TurboID-TTC5 fusion reconstitutes autoregulation in a TTC5 knockout (KO) cell line and that the TurboID-TTC5(K97A) mutant is ineffective (Fig. S1E), we induced biotinylation during active autoregulation (Fig. S1F) and affinity-purified the biotinylated proteins (Fig. S1G). Quantitative mass spectrometry revealed that a poorly studied protein named SCAPER (S-phase Cyclin A Associated Protein residing in the ER) was the only protein whose biotinylation was strongly reduced in TTC5(K97A) cells (Fig. 1C). Notwithstanding its name, SCAPER lacks obvious ER-targeting domains and is nucleo-cytoplasmic as determined by immunostaining^36^. Immunoblotting verified that in cells, SCAPER is biotinylated by TurboID-TTC5 in a K97-dependent manner (Fig. 1D).

In pulldown experiments, purified SCAPER interacted with purified TTC5 but not TTC5(K97A) (Fig. 2A). Structure modelling using AlphaFold2 (AF2) multimer^37,38^ predicted a high-confidence interaction between the region of TTC5 containing K97 and a globular C-terminal domain (CTD) of SCAPER (Fig. S2A and S2B). In a cytosolic in vitro translation reaction, recombinant SCAPER co-fractionated and co-purified with TTC5-RNC complexes displaying the first 64 amino acids of β-tubulin (Fig. 2B and 2C; Fig. S2C). This interaction was not seen in reactions containing TTC5(K97A), reactions lacking β-tubulin RNCs, or reactions containing RNCs with mutant β-tubulin incapable of TTC5 recruitment. Thus, SCAPER is selectively recruited to tubulin-synthesizing ribosomes via a direct interaction with TTC5.

**Figure 2.**
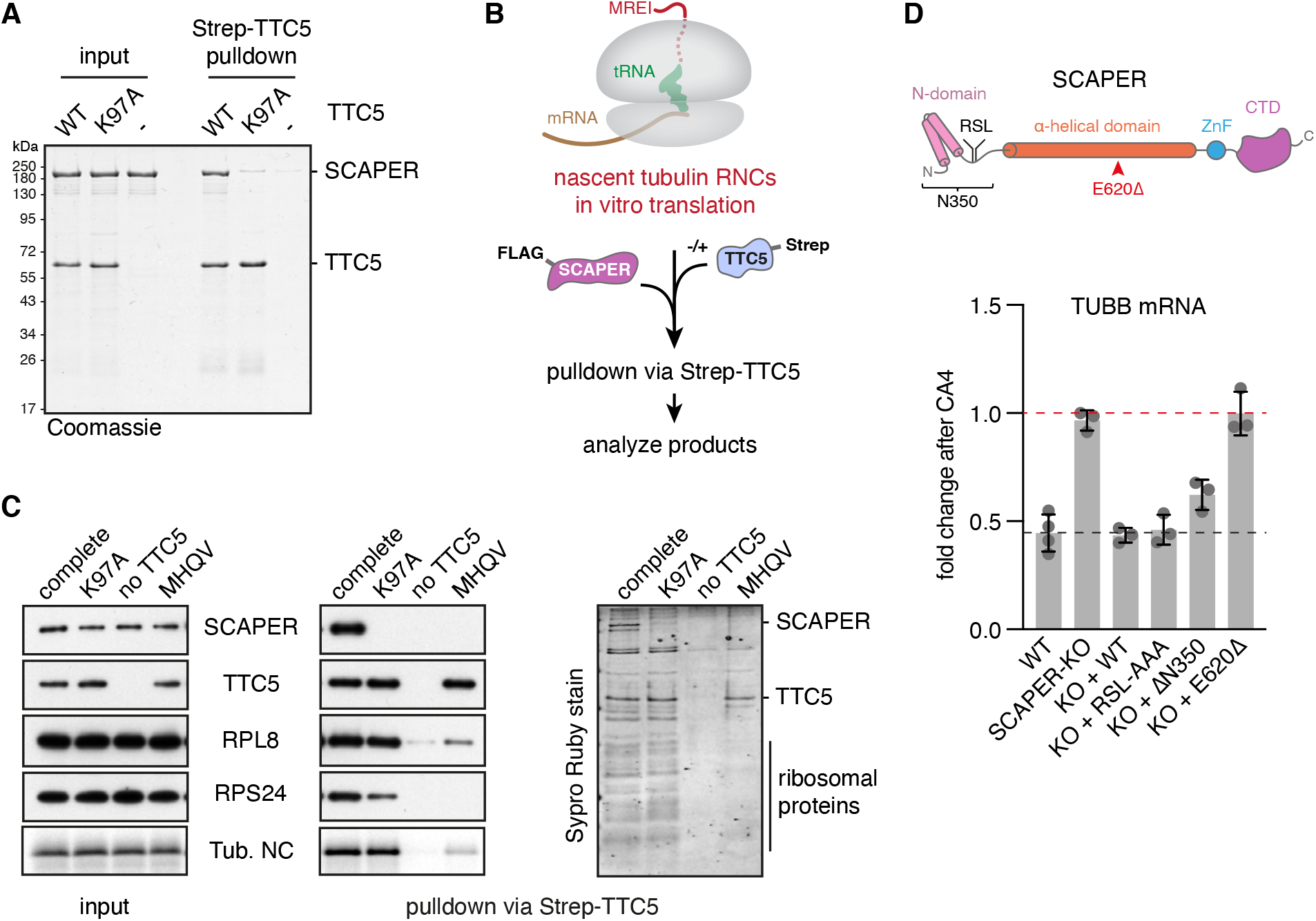
TTC5 recruits SCAPER to tubulin ribosome nascent chain complexes for autoregulation. (A) Recombinant Strep-TTC5 and SCAPER-FLAG were incubated together and pulled down via Strep-TTC5. Bound proteins were separated using SDS-PAGE and visualized by Coomassie staining. (B) Schematic workflow for reconstitution of SCAPER recruitment to tubulin ribosome nascent chains (RNCs) via TTC5 as shown in panel C. (C) 64-residue β-tubulin (TUBB) nascent chains were produced in rabbit reticulocyte lysates in the presence of recombinant FLAG-SCAPER (all samples) and Strep-TTC5 as indicated, and TTC5-associated proteins were subsequently enriched via its Strep tag. Input and Strep-TTC5 pulldown samples were separated by SDS-PAGE and visualized by western blotting, autoradiography for the β-tubulin nascent chain (Tub. NC), or SYPRO Ruby staining for total protein. “MHQV” indicates a β-tubulin construct in which its TTC5-interacting MREI motif has been mutated. (D) Top panel: Schematic of SCAPER domain architecture including annotated features and predicted structural elements. The pathologic E620Δ mutation is indicated by a red arrow head (see also Fig. S4A). RSL: cyclin A binding motif (Arg^199^-Ser^200^-Leu^201^); ZnF: Zinc finger; CTD: Carboxy-terminal domain. Bottom panel: Autoregulation assay with wild type, SCAPER-KO and the indicated rescue cell lines. RSL-AAA: mutation of the cyclin A binding site (Arg^199^-Ser^200^-Leu^201^) to alanines; N350: deletion of the first 350 residues upstream of the α-helical domain; E620Δ: deletion of residue Glu^620^. Mean +/− SD from 3 replicates is plotted.

### SCAPER is required for autoregulation

Cells knocked down or knocked out for SCAPER are completely deficient in tubulin autoregulation (Fig. S3) and can be fully rescued by SCAPER reintroduction (Fig. 2D; Fig. S3E and S3F). Domain mapping experiments showed that the N-terminal part, which contains the SCAPER N-domain and a previously characterized cyclin A binding site^39^, is largely dispensable for autoregulation (ΔN350 in Fig. 2D; Fig. S4A–C). Consistent with this result, a cyclin A binding mutant (RSL-AAA) had no effect on autoregulation, whereas the central α-helical domain and CTD were required (Fig. 2D; Fig. S4B and S4C).

Interestingly, numerous disease-linked SCAPER mutations cause C-terminal truncations or are located in the central and C-terminal domains (Fig. S4A)^40,41^. These mutations lead to retinitis pigmentosa, intellectual disability, male infertility and other pathologies consistent with microtubule cytoskeleton aberrations^41–43^. None of the truncation mutants are expected to be competent for autoregulation given the crucial role of SCAPER’s CTD.

Furthermore, two disease-causing deletion mutants in the central α-helical domain (E620Δ and 675-677Δ) led to severe autoregulation defects without appreciably affecting SCAPER expression (Fig. 2D; Fig. S4D and S4E), whereas a third disease allele (S1219N) was expressed at substantially lower levels (Fig. S4E), presumably due to destabilization of the protein. Notably, the E620A substitution mutant was fully functional, hinting that the α-helix register in this part of SCAPER might be crucial for its function (Fig. S4D and S4E). Thus, SCAPER alleles that cause human disease are impaired in tubulin autoregulation, highlighting key roles for the autoregulation pathway in human physiology.

### Mechanism of ribosome engagement by SCAPER

To understand how SCAPER binds tubulin-synthesizing ribosomes, we analyzed β-tubulin-RNCs engaged with recombinant TTC5 and SCAPER (Fig. 2C) by single-particle cryo-electron microscopy (cryo-EM). The structure, at an overall resolution of 2.8 Å and local resolution from 3 to 8 Å for non-ribosomal regions (Supplementary Table S1), showed that SCAPER’s CTD makes contacts with TTC5, the 60S surface, and an additional density that was identified as the 28S rRNA expansion segment ES27L (Fig. 3A; Fig. S5). The other parts of SCAPER upstream of residue 859 were not resolved. AF2 models of TTC5 with the tubulin nascent chain and the SCAPER-CTD were docked into the cryo-EM map and adjusted to generate a structural model.

**Figure 3.**
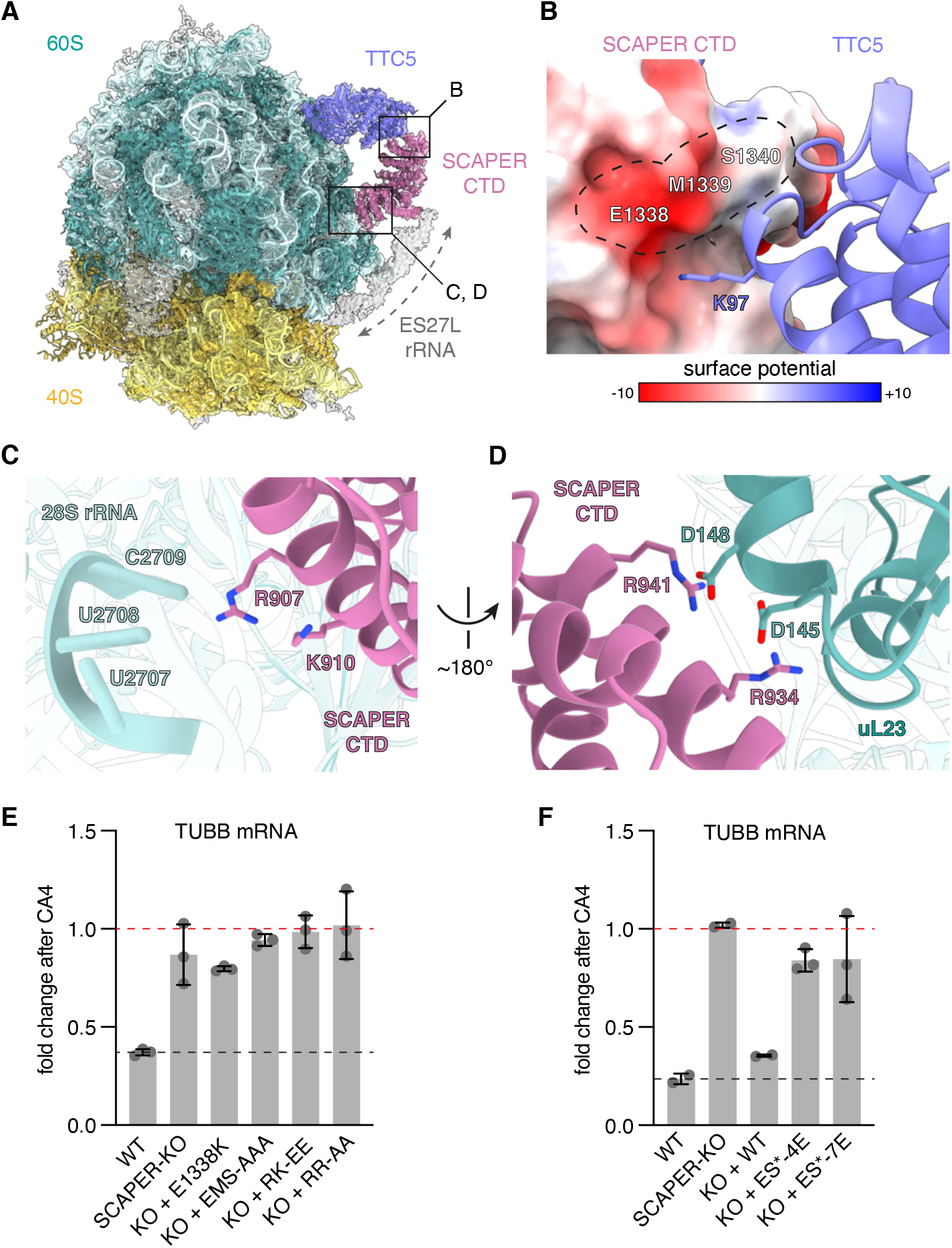
Mechanism of SCAPER recruitment to tubulin RNCs via TTC5. (A) Overview of the cryo-EM-derived structure of β-tubulin-synthesizing ribosomes bound to TTC5 and SCAPER. Dashed arrow marks density that was identified as 28S rRNA expansion segment ES27L. Boxes indicate positions of close-ups shown in panels B–D. The displayed non-sharpened map resulted from the ES27L classification (see Fig. S5). The 40S subunit was rigid-body docked and is shown to orient the reader. (B) Close-up view of the contact between SCAPER and TTC5. SCAPER is coloured by electrostatic surface potential [in kcal/(mol\**e*)], and the surface area of critical residues is outlined. (C) Close-up view of critical positively charged SCAPER residues in close vicinity to 28S rRNA. (D) Close-up view of conserved arginine residues of SCAPER in close proximity to aspartate residues of ribosomal protein uL23. (E) Autoregulation assay comparing WT, SCAPER-KO and rescue cell lines expressing the indicated SCAPER mutants. EMS-AAA: E1338A, M1339A, S1340A; RK-EE: R907E, K910E; RR-AA: R934A, R941A. (F) Autoregulation assay as in panel E with mutations in the expansion segment contact site of SCAPER. ES*-4E: K867E, K870E, K873E and K874E; ES*-7E: as ES*-4E plus K869E, K871E and R878E. See also Fig. S6. Bar graphs show mean +/− SD.

In this model, K97 of TTC5 is positioned near a negatively charged and highly conserved surface patch on SCAPER around E1338, explaining why the K97A mutation is defective in SCAPER interaction and autoregulation (Fig. 3B). Furthermore, two conserved positively charged surface patches on SCAPER contact the 60S subunit and ES27L (Fig. S6A). At the 60S interface, R907 and K910 of SCAPER abut 28S rRNA residues U2707–C2709, and R934 and R941 of SCAPER interact with D145 and D148 of ribosomal protein uL23 (Fig. 3C and 3D, respectively). At the ES27L interface, a cluster of eight conserved positively charged residues between K867 and R878 along an α-helix from SCAPER faces rRNA (Fig. S6B), although details of this interaction were not visualized at the moderate resolution in this part of the map. The function of rRNA expansion segments is poorly understood, but ES27L emerges as a key structural element that is known to scaffold binding of factors around the exit tunnel for various functions^44–46^.

SCAPER variants with point mutations at the interaction sites with TTC5, the 60S body, and ES27L each failed to restore autoregulation to SCAPER KO cells (Fig. 3E and 3F) despite high expression levels (Fig. S6C and S6D). The charge reversal mutation E1338K in SCAPER, opposite to K97 in TTC5, strongly impacted autoregulation. Similarly, a triple alanine mutation of E1338, M1339, and S1340 (EMS-AAA) on the SCAPER surface that forms the primary TTC5 binding site was completely inactive. Finally, mutants of conserved positively charged SCAPER residues that contact either the 60S body rRNA (R907E, K910E), uL23 (R934A, R941A), or ES27L (ES*-4E or -7E) were inactive. Thus, SCAPER uses its CTD to selectively engage TTC5-containing ribosomes through three crucial contacts. The structure explains why all disease-causing premature termination codons in SCAPER (Fig. S4A), even those close to the C-terminus, would be incompatible with SCAPER recruitment by TTC5. Furthermore, the region N-terminal to the ribosome-binding CTD would extend toward the 40S subunit and potentially reach over 300 Å (Fig. S4A). This is noteworthy because SCAPER can readily bridge the distance from the polypeptide exit tunnel, where TTC5 binds the nascent chain, to the 40S subunit through which the mRNA is threaded.

### SCAPER recruits CCR4-NOT for mRNA deadenylation

Because SCAPER has no apparent catalytic domains that would degrade mRNA, we speculated it acts as an adaptor that recruits a nuclease. The absence of a nuclease in our TTC5-centered proximity labelling experiment hinted that distal regions of SCAPER too far for proximity biotinylation might mediate nuclease recruitment. We therefore repeated the experiment with TurboID fused to the N-terminus of SCAPER (Fig. 4A). The set of biotinylated proteins recovered from cells that are acutely degrading tubulin mRNA, relative to cells at steady state, was enriched for multiple subunits of the CCR4-NOT deadenylase complex (Fig. S7A). Strikingly, biotinylated CCR4-NOT subunits were strongly de-enriched in samples from cells expressing SCAPER(E620Δ), a pathologic mutant defective in tubulin autoregulation (Fig. 4B; Fig. 2D). Thus, CCR4-NOT is proximal to SCAPER’s N-terminus preferentially during autoregulation conditions of active tubulin mRNA degradation.

**Figure 4.**
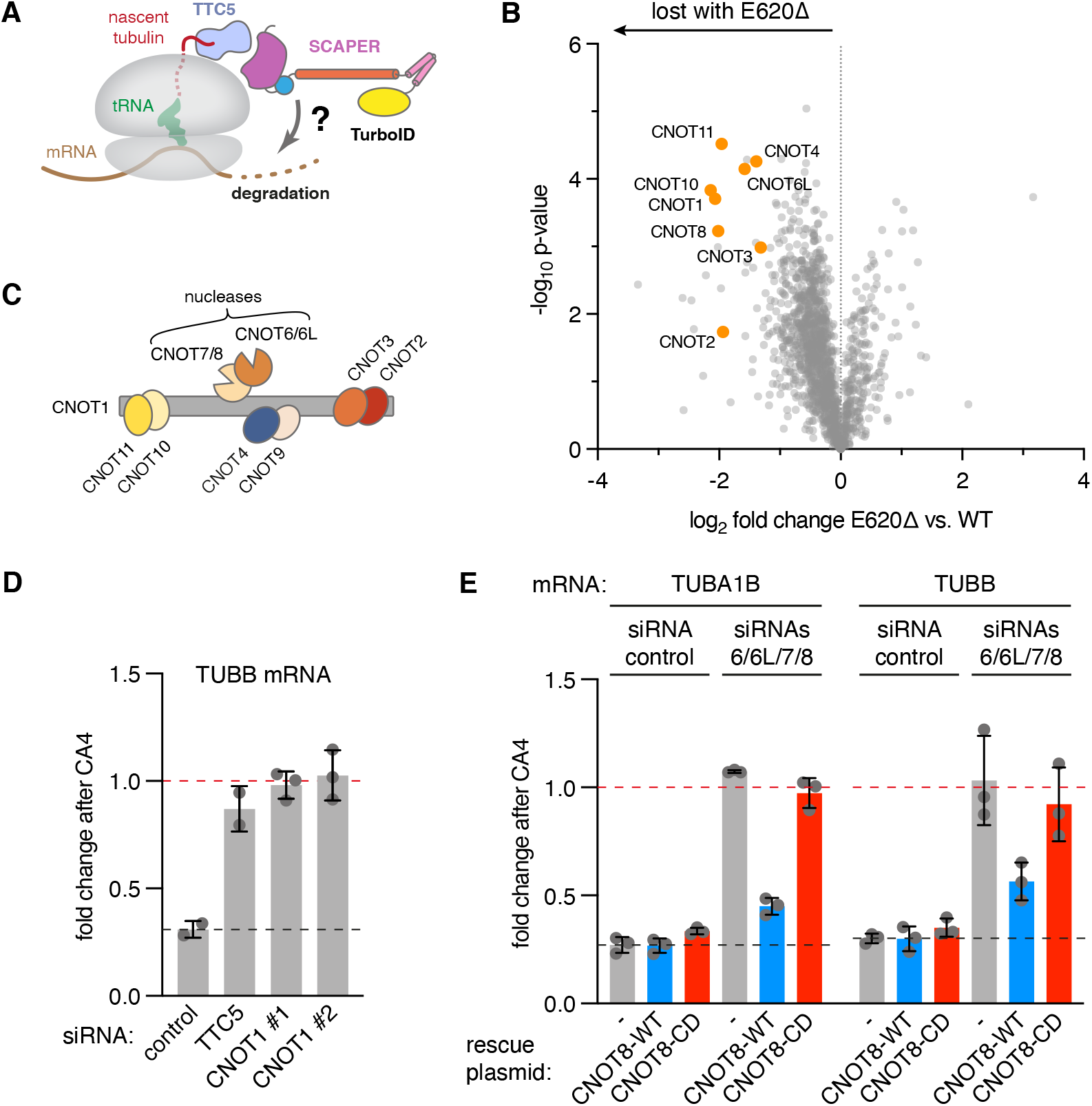
The CCR4-NOT complex triggers tubulin mRNA degradation. (A) Schematic for the strategy to identify factors acting downstream of SCAPER to degrade tubulin mRNA. (B) Proximity labelling using TurboID fusions to either SCAPER WT or the E620Δ mutant, performed after induction of autoregulation with the microtubule depolymerization agent Combretastatin A4 (CA4, at 100 nM). Biotinylated proteins were enriched and analyzed by quantitative mass spectrometry. See also Supplementary Table S3. (C) Schematic of the subunit composition of the CCR4-NOT complex. (D) Autoregulation assay performed after knockdowns (KD) using control, TTC5- or CNOT1-targeted siRNAs. (E) Autoregulation assays performed after control KD, or KD of all partially redundant catalytic deadenylase subunits of CCR4-NOT (siRNAs 6/6L/7/8). CNOT8 expression was accomplished by stable integration of siRNA-resistant CNOT8-WT (blue bars) or the catalytic dead D40A mutant (CNOT8-CD, red bars). Bar graphs show mean +/− SD.

The CCR4-NOT complex is a large multi-subunit complex (Fig. 4C) responsible for most cytoplasmic deadenylation activity, the first and often rate-limiting step in mRNA decay^22,47,48^. siRNA-mediated knockdown (KD) of CNOT1, the large scaffolding subunit around which all other subunits assemble, completely abolished tubulin mRNA degradation in autoregulation assays (Fig. 4D; Fig. S7B). Similarly, knockdown of all four partially redundant nuclease subunits (CNOT6, CNOT6L, CNOT7, CNOT8) also stabilized tubulin mRNAs (Fig. 4E; Fig. S7C). This defect was rescued by re-expressing siRNA-resistant CNOT8, but not by a catalytic dead mutant (Fig. 4E; Fig. S7C). Thus, the CCR4-NOT complex is physically proximal to SCAPER during autoregulation and its deadenylase activity is required for tubulin mRNA degradation.

### Mechanism of CCR4-NOT recruitment by SCAPER

CNOT1 not only scaffolds the catalytic exonuclease subunits, but also regulatory subunits that deploy the CCR4-NOT complex to specific mRNAs via RNA-binding adaptor proteins^22,48^. An initial screen of all CCR4-NOT subunits by siRNA-mediated KD revealed that CNOT10 and CNOT11 are most important for tubulin autoregulation (Fig. 5A; Fig. S7D). The specificity of this effect was underscored by the finding that several other previously described CCR4-NOT substrates were stabilized by CNOT1 KD but not by KD of CNOT10 or CNOT11 (Fig. 5B), consistent with previous findings^49^. CNOT10 interacts with CNOT11 to form a module that evolved later than the core CCR4-NOT complex, similar to the evolution of other tubulin autoregulation components^16,50^. This suggested that the CNOT10/CNOT11 module, while dispensable for some other CCR4-NOT functions, might recognize SCAPER for recruitment to tubulin RNCs. Supporting the notion of a specific role in tubulin homeostasis, a recent study found steady-state tubulin protein levels elevated upon CCR4-NOT disruption, specifically when CNOT1, CNOT10 or CNOT11 were depleted^51^.

**Figure 5.**
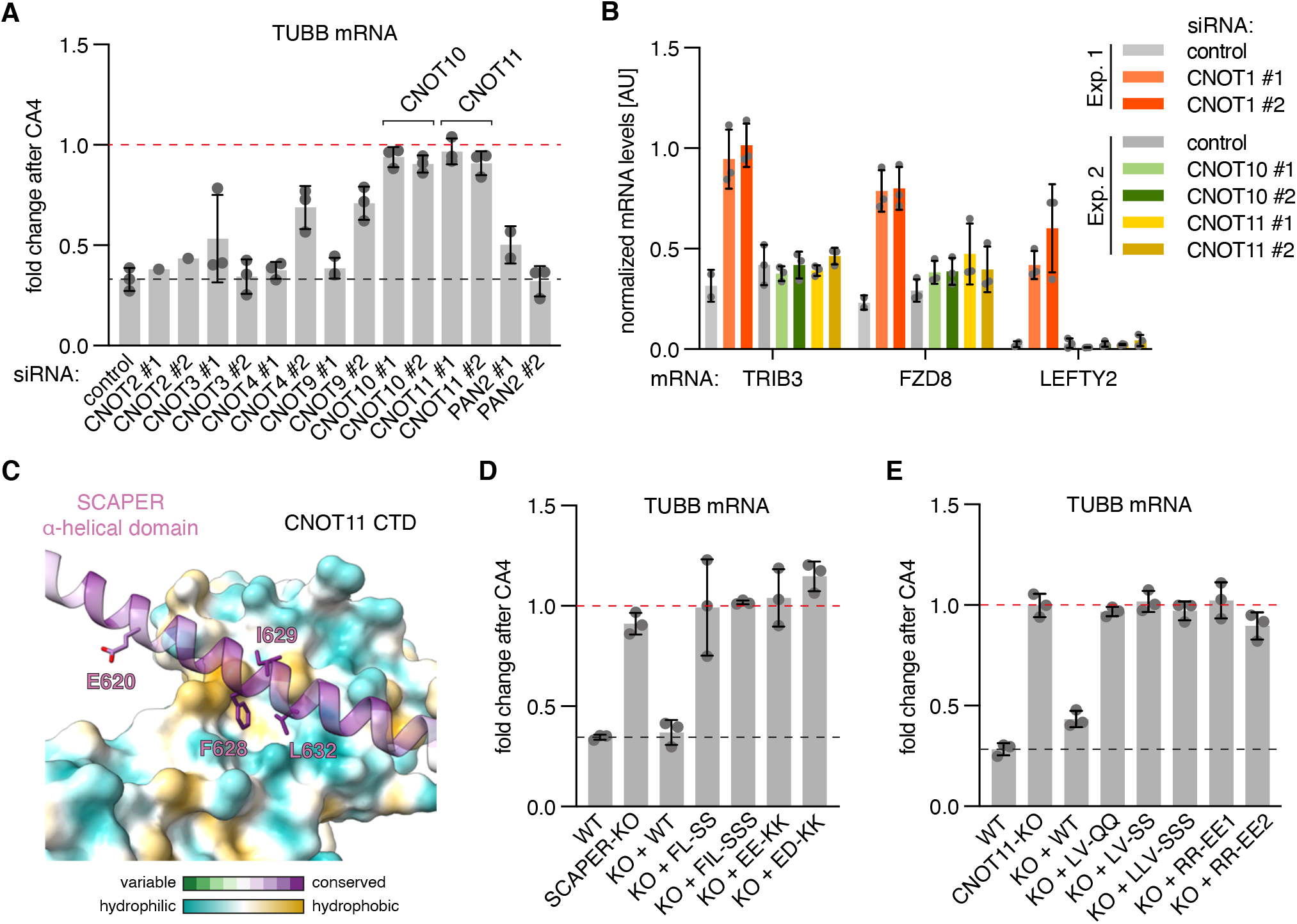
SCAPER recruits the CCR4-NOT complex via CNOT11. (A) Autoregulation assays were performed after knockdown using the indicated siRNAs for 3–4 days. Data are from one experiment for CNOT2-KD, and mean +/− SD from 2–3 replicates for all other siRNAs. We note that KD of PAN2 did not lead to stabilization of tubulin mRNAs in autoregulation assays. PAN2 is the catalytic subunit of the PAN2-PAN3 complex that often initiates deadenylation prior to CCR4-NOT^22^. (B) RT-qPCR quantification of previously identified CCR4-NOT substrates^66,67^ in samples with KD for CNOT1, CNOT10, or CNOT11. The same samples from control conditions used in Fig. 4D (Exp. 1) and Figure 5A (Exp. 2) were analyzed. Target mRNA levels were normalized to a house-keeping gene (GAPDH). Normalization to 18S rRNA, which is not a deadenylation substrate of CCR4-NOT, gave comparable results. Note that LEFTY2 mRNA levels were at or below the detection threshold for all samples except CNOT1-KD samples. (C) Model of AlphaFold2 multimer predicted interaction between the C-terminal domain of CNOT11 with the α-helical domain of SCAPER. E620 and three highly conserved hydrophobic SCAPER residues predicted to interact with a hydrophobic patch on CNOT11 are highlighted. (D) Autoregulation assay with WT, SCAPER-KO, or SCAPER rescue cell lines with the indicated mutations targeting the predicted CNOT11 interaction surface. FL-SS: F628S, L632S; FIL-SSS: F628S, I629K, L632S; EE-KK: E618K, E625K; ED-KK: E633K, D640K (E) Autoregulation assay with WT, CNOT11-KO, or CNOT11 rescue cell lines with the indicated mutations targeting the predicted SCAPER interaction surface. LV-QQ: L405Q, V454Q; LV-SS: L405S, V454S; LLV-SSS: L405S, L451S, V454S; RR-EE1: R447E, R450E; RR-EE2: R461E, R485E. Bar graphs show mean +/− SD.

Other subunits of CCR4-NOT, such as CNOT2, CNOT3, and CNOT9 interact with substrate-specific RNA binding proteins that act as adaptors for selective mRNA decay. Speculating that SCAPER might be a substrate-specific adaptor for the CNOT10/CNOT11 module, we screened for potential interactions with regions of SCAPER using AF2 multimer^37,38^. A high-confidence interaction was predicted between the highly conserved C-terminal DUF2363 domain of CNOT11 and conserved residues of the SCAPER α-helical domain (Fig. 5C and Fig. S8). Strikingly, E620 of SCAPER was adjacent to the CNOT11 binding surface (but not in direct contact), perhaps explaining why a shift of α-helix register in this region caused by the E620Δ mutation abolishes autoregulation and causes disease (Fig. 5C; Fig. S8C; Fig. 2D).

Guided by the AF2 prediction, we designed mutations on either side of the SCAPER-CNOT11 interface and introduced them into the respective knockout cell line. Whereas the wild type constructs rescued the knockout phenotype, each of the interface mutants was completely deficient for autoregulation (Fig. 5D and 5E; Fig. S8C and S8D; Fig. S7E and S7F), validating key features of the AF2-predicted interaction. Taken together, our data imply that the CCR4-NOT complex employs its CNOT10/CNOT11 module to selectively engage tubulin RNCs marked by the TTC5-SCAPER complex via nascent chain recognition. At these RNCs, the nuclease subunits of CCR4-NOT can deadenylate tubulin mRNAs to trigger their degradation during autoregulation.

### SCAPER mutation causes mitosis defects

Accurate regulation of tubulin levels is crucial for MT-dependent processes, including the formation of the mitotic spindle during cell division. To investigate the relevance of SCAPER-dependent autoregulation during mitosis in a cell-based assay, we monitored chromosome segregation using live cell microscopy (Fig. 6A). We found that SCAPER KO cells have a ~4-fold increase in chromosome alignment and segregation errors (Fig. 6B and 6C; Fig. S9) similar to the effects seen in TTC5 KO cells^17^. Neither the cyclin A binding site mutation (RSL-AAA) nor truncation of the N-terminus showed this phenotype. In contrast, the E620Δ disease mutant, which is deficient in CCR4-NOT recruitment, essentially phenocopied the SCAPER KO (Fig. 6B and 6C; Fig. S9). These outcomes closely correlate with the phenotypes in tubulin autoregulation assays of the respective genotypes (Fig. 2D, Fig. S9B). The observed mitosis defects are expected to result in aneuploidy, which is associated with cancer progression and can impair neurodevelopment^52–54^. These data indicate that SCAPER is required for faithful mitosis, most likely through fine-tuning of microtubule dynamics via selective tubulin mRNA degradation.

**Figure 6.**
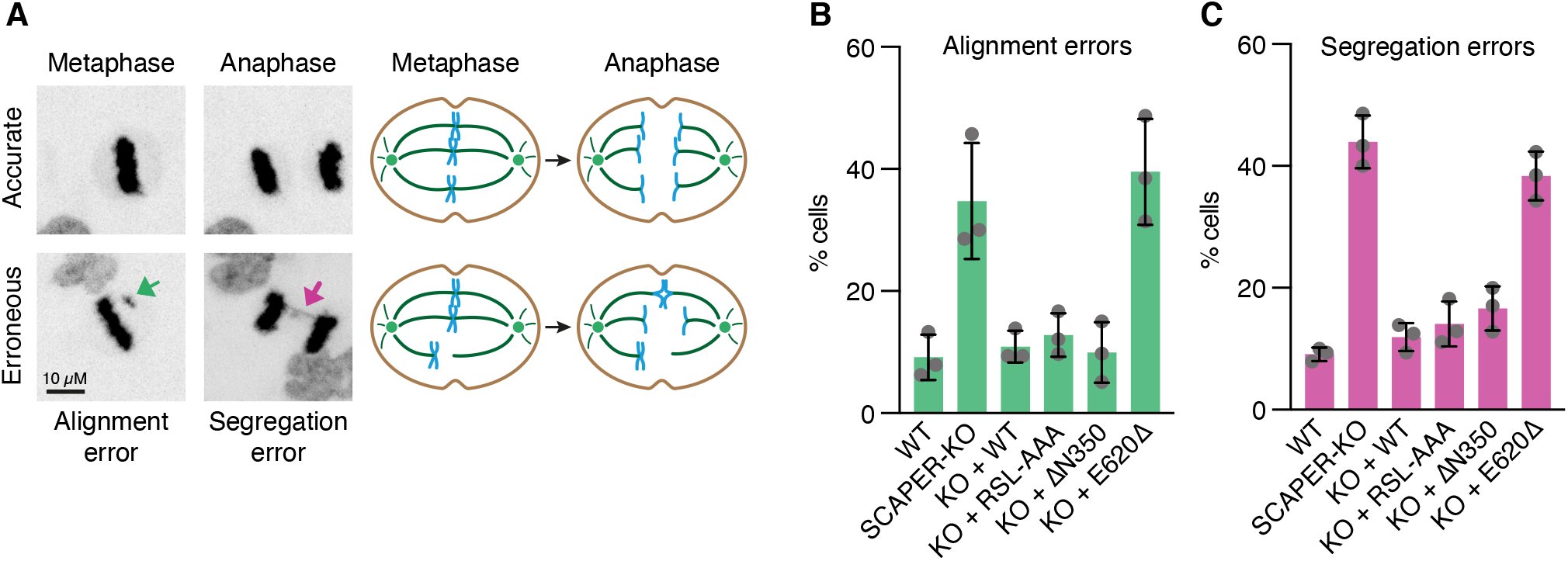
SCAPER is required for accurate mitosis. (A) Left panels: Example images of meta- and anaphase stages of cells going through mitosis where chromosomes were visualized using SiR-DNA stain and maximum intensity projections are shown. Misaligned chromosomes and segregation errors are highlighted by green and magenta arrows, respectively. Right panels: Schematics of accurate (top) and erroneous (bottom) cell division stages. Chromosomes are shown in blue, MTs in dark green, centrosomes in light green. (B) Quantification of chromosome alignment errors in stable Flp-In HeLa T-REx cell lines with the indicated genotypes. (C) Quantification of chromosome segregation errors in stable Flp-In HeLa T-REx cell lines with the indicated genotypes. Data show mean +/− SD from three biological replicates with 100 cells in total for (B) and (C).

## DISCUSSION

The mechanistic basis for selective tubulin mRNA degradation and its physiological function have been long-standing questions since the description of tubulin autoregulation more than forty years ago^14^. In this work, we elucidated the factors and interactions that bridge nascent tubulin peptide recognition at the ribosome exit tunnel to mRNA deadenylation (Fig. 7). The findings assign molecular functions to the previously obscure proteins SCAPER and CNOT11, provide mechanistic insight into genetic diseases caused by SCAPER mutations, and provide the first detailed view of how a nascent protein can selectively control the degradation of its encoding mRNA. The work therefore highlights new principles in post-transcriptional gene regulation.

**Figure 7.**
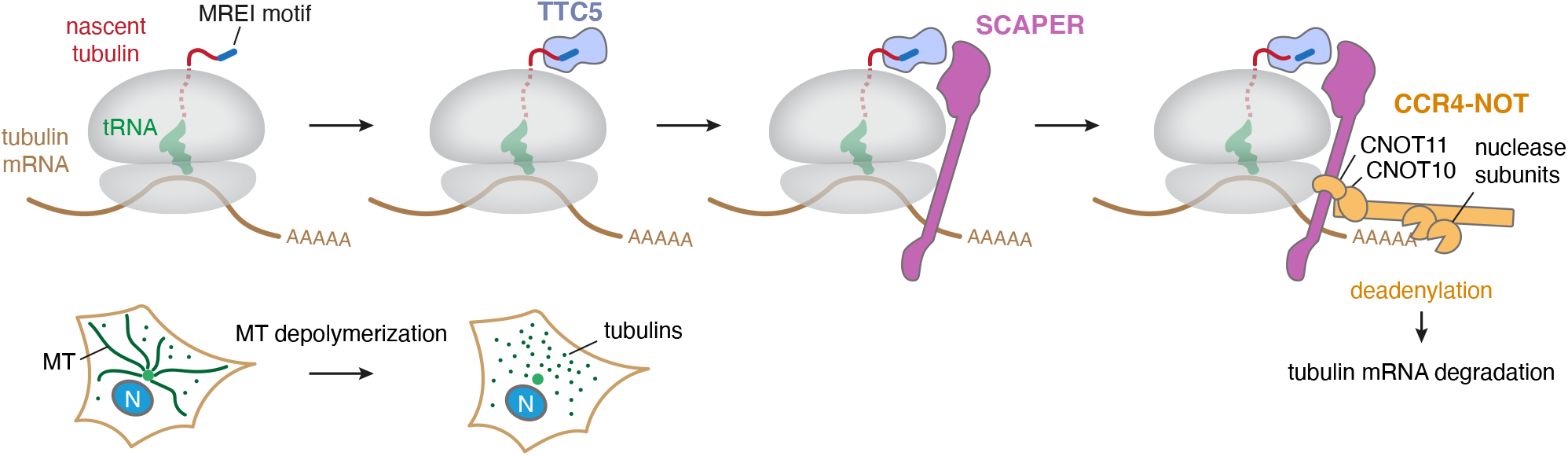
Model of regulated mRNA degradation in the tubulin autoregulation pathway. Selective tubulin mRNA degradation is triggered when cells sense excess free tubulin levels, e.g. due to microtubule (MT) depolymerization, as depicted in the bottom schematic (N: nucleus). Under these conditions TTC5 is liberated from an elusive inhibitory factor^17^ (not shown). This allows TTC5 to selectively bind tubulin-translating ribosomes by interacting with the conserved N-terminal peptide motif (Met-Arg-Glu-Ile or MREI, shown in dark blue) and a surface around the ribosomal exit tunnel. SCAPER recruitment is in turn facilitated by a composite interaction surface formed by TTC5 and the ribosome. The CCR4-NOT complex uses its CNOT11 subunit to bind an extended α-helical domain of SCAPER and its nuclease subunit(s) to deadenylate tubulin mRNA to initiate its subsequent degradation.

The most noteworthy insight to emerge from our studies is the mechanistic basis for how an mRNA can be targeted for selective degradation by direct recognition of the nascent protein. Instead of sequence-specific recognition of tubulin mRNAs, a series of protein-protein interactions at the translating ribosome culminates in recruitment of a general deadenylase complex. A major advantage of this mechanism is that an entire class of mRNAs, the α- and β-tubulins totalling 19 genes in humans, can be targeted as a group despite widely varying UTRs and coding sequences. Instead, they are recognized via a shared peptide motif in the proteins they encode. This is conceptually analogous to how a single microRNA can coordinately regulate multiple widely different proteins based on a shared recognition motif in their encoding mRNAs^21^.

In the autoregulation pathway, TTC5 imparts specificity for tubulins and contributes decisively to the specificity of SCAPER recruitment. Because SCAPER has the potential to be highly elongated, the CNOT11 binding site can reach far from the polypeptide exit tunnel where its CTD engages TTC5. Consistent with this idea, crosslinking mass spectrometry experiments suggest that CNOT11 may contact the mRNA-binding 40S subunit^55^. SCAPER therefore acts as a molecular bridge that effectively communicates a nascent chain recognition event at the exit tunnel on the ribosome 60S subunit to a deadenylase activity that likely resides near the mRNA channel of the 40S subunit. Thus, CCR4-NOT can be deployed to selective ribosomes on the basis of the nascent polypeptides they display, a new mode of action qualitatively different from direct binding to either ribosomes or sequence-specific mRNA-binding adaptors^22,48,56^.

Our mechanistic dissection of nascent tubulin-dependent recruitment of the CCR4-NOT complex provides a framework for understanding analogous regulatory processes for other proteins. For example, the stability of mRNAs coding for at least some ribosomal proteins is coupled to the availability of chaperones dedicated to these proteins in budding yeast^25^. Degradation of these mRNAs in the absence of chaperones is thought to be co-translational, but neither the basis of nascent chain recognition nor the mechanism of putative CCR4-NOT recruitment are understood. The methods and principles from the tubulin autoregulation pathway provide a roadmap to now dissect the analogous processes for ribosomal proteins and others.

Mutations in TTC5 and the N-terminal recognition motif in a tubulin gene have previously been linked to neurodevelopmental defects^57,58^, hinting at physiologic role(s) for autoregulation. However, potential added roles for TTC5 in regulation of transcription and the actin cytoskeleton^59,60^, and putative consequences for tubulin structure complicated this interpretation. Our assignment of SCAPER to the autoregulation pathway and characterization of autoregulation-disrupting mutants now strengthen substantially the link between autoregulation and human physiology. SCAPER disease variants lead to ciliopathy-related syndromes comprising intellectual disability, retinitis pigmentosa, male infertility, and other symptoms^41–43^. These phenotypes overlap partially with both TTC5-linked disease and tubulinopathies, providing insights into the tissues and biological processes most reliant on tubulin autoregulation. Interestingly, the nervous system is exquisitely sensitive to mutations that cause DNA damage or chromosome segregation defects, possibly due to the rapid proliferation of neuronal progenitor cells required during brain development^53,54,61^. Thus, the complex phenotypes seen in humans mutant for SCAPER may be due to a combination of defective ciliogenesis, chromosome segregation, and some of the many other tubulin-related processes.

How the tubulin autoregulation pathway is controlled in response to changes in microtubule or free tubulin levels remains enigmatic. In this regard, the previously identified cyclin A binding site of SCAPER^39^, and its putative microtubule binding activity^43^ suggest potential mechanisms of regulation. Indeed, tubulin mRNA levels have been observed to change through the cell cycle as might be needed to accommodate different roles of the microtubule network^62,63^. More generally, multi-component pathways provide ample scope for temporal and context-dependent regulation^64,65^. In tubulin autoregulation, the specificity factor TTC5, the adaptor SCAPER, the substrate-recruitment subunit CNOT11, and the deadenylase complex CCR4-NOT could all be fine-tuned to ensure accurate tubulin levels in a cell type-specific manner. Our work establishes a framework to study other co-translational mechanisms that use nascent chain features to direct diverse mRNA fates.

## Acknowledgments

We are grateful to the electron microscopy facility staff at the MRC Laboratory of Molecular Biology for access and support for sample preparation and data collection; J. Grimmett and T. Darling for maintenance of the LMB scientific computing infrastructure; F. Begum and M. Skehel for mass spectrometry; C. Lau and S. Chaaban for help with a local implementation of AlphaFold2; V. Chandrasekaran and J. P. O’Donnell for comments on the manuscript; the Photonic Bioimaging Center of the University of Geneva for help with microscopy. This work was supported by the Medical Research Council, as part of United Kingdom Research and Innovation (MC_U105192715 to L.A.P. and MC_UP_A022_1007 to R.S.H.). M.H. received funding from EMBO (ALTF 116-2020) and the European Union’s Horizon 2020 research and innovation programme under the Marie Sklodowska-Curie grant agreement No 101029853. E.A. was funded by a postdoctoral fellowship of the Deutsche Forschungsgemeinschaft (DFG, German Research Foundation). I.G. is supported by the Swiss National Science Foundation Eccellenza Fellowship (PCEFP3_194312) and is the Dale F. Frey Breakthrough Scientist of the Damon Runyon Cancer Research Foundation (DRG:2279-16). We acknowledge Diamond Light Source for access to eBIC (proposal BI23268) funded by the Wellcome Trust, Medical Research Council and Biotechnology and Biological Sciences Research Council.

## Author Contributions

M.H. discovered SCAPER and performed most experiments; E.A. performed structural analysis of the TTC5-SCAPER-ribosome complex; S.Y.P.C. analysed mass spectrometry samples; I.G. and E.V. generated and validated HeLa cell lines; I.G. designed and performed phenotypic analysis of mitosis. R.S.H., L.A.P. and I.G. supervised different parts of the project. R.S.H. and M.H. conceived the project, oversaw its implementation, and wrote the manuscript. All authors contributed to manuscript editing.

## METHODS

### Plasmids and reagents

β-tubulin (human TUBB) constructs for in vitro translation have been described previously^17^. EGFP-tagged TTC5 (“GFP-TTC5”) was obtained by cloning previously described TTC5 constructs^17^ into a pcDNA5/FRT/TO with an N-terminal EGFP tag. N-terminally 6xHis-TEV-Twin-Strep-tagged TTC5 (“Strep-TTC5”) for bacterial expression was cloned in the pET-28a vector. A human SCAPER cDNA construct with C-terminal FLAG-tag in a pcDNA3.1 vector was obtained from Genscript (cloneID OHu03552) and subsequently cloned into pcDNA5/FRT/TO vectors with N- or C-terminal FLAG-tags. TurboID-FLAG was fused to the N-terminus of TTC5 or SCAPER (WT or mutants) and cloned into pcDNA5/FRT/TO vectors. Human siRNA-resistant CNOT8-WT and -CD were cloned from synthetic gene blocks (IDT) into pcDNA5/FRT/TO with a C-terminal PreScission cleavage site followed by a Twin-Strep-tag. Human CNOT11 was cloned from HEK293 T-REx cDNA into pcDNA5/FRT/TO with an N-terminal 3HA-TEV-tag. CRISPick (https://portals.broadinstitute.org/gppx/crispick/public) was used to design sgRNAs for CRISPR-Cas9-mediated knockout (KO) of SCAPER and CNOT11. The sequences were as follows:

SCAPER-KO sgRNA #1 (exon 5): TTGCGAATCAGATCAGAGTG
SCAPER-KO sgRNA #3 (exon 17): AAGCCCGAATTGAACAACAG
CNOT11-KO sgRNA AC (exon 2): CAACGTCCATCAGTGCAATC

### Cell culture procedures

HEK 293 Flp-In T-REx cells (Invitrogen) were maintained in DMEM with GlutaMAX and 4.5 g/l glucose (Gibco) supplemented with 10% fetal calf serum, and optionally 0.1 mg/ml Hygromycin B and 10 μg/ml Blasticidine S for stable Flp-In cell lines. CRISPR-Cas9 mediated gene knockout for SCAPER was performed essentially as described^68^: HeLa or HEK293 Flp-In TRex cells were transiently transfected with the pX459 plasmid encoding the sgRNAs targeting SCAPER and Cas9, using Lipofectamine 3000 reagent (Invitrogen) for HeLa cells or TransIT-293 (Mirus) for HEK T-REx cells following manufacturers’ protocols. 24 hours after transfection, 2 μg/ml puromycin (1μg/ml for HEK293) was added for selection. 2–3 days after transfection, cells were trypsinized and re-plated in 96-well plates at a density of 0.5 or 1 cell per well using a FACSAria Fusion instrument (BD) to obtain single cell clones. To obtain CNOT11-KO clones, IDT Alt-R sgRNA was complexed with Alt-R S.p. Cas9 Nuclease V3 and transfected into HEK T-REx cells using Lipofectamine RNAiMAX (Invitrogen) according to the IDT user guide. Cells were grown for 48 hours and then sorted into 96-well plates.

Successful knockout clones were verified by genotyping via PCR amplification of the modified region followed by TIDE analysis^69^. CNOT11-Kos were further confirmed by western blotting, but no suitable antibody for detection of endogenous SCAPER levels could be sourced. Rescue cell lines with stable expression of TTC5, SCAPER, CNOT8 or CNOT11 constructs were generated in knockout cells (or wild type cells for CNOT8) using the Flp-In system (Invitrogen) following manufacturer’s protocol. Expression of transgenes was induced with 200 ng/mL (HeLa) or 1 μg/ml (HEK T-REx) doxycycline for 24–48 hours. Colchicine (10 *μ*M), Nocodazole (10 *μ*M), and combretastatin A4 (CA4, 100 nM) treatments were performed in standard media for 3 h, unless stated otherwise. All drugs gave similar effects in autoregulation assays, but we found results with colchicine more variable and hence used CA4 throughout most of the study, which gave consistent results.

For siRNA mediated knockdowns of indicated genes, Silencer Select siRNAs (Thermo Fisher) were transfected using RNAiMAX (Invitrogen) according the manufacturer’s instructions for reverse transfection. Cells were typically incubated for three days, unless stated otherwise. When multiple siRNAs were transfected, they were used in equal ratios with the total amount of siRNA kept constant.

### Live cell imaging and data analysis

Flp-In T-REx HeLa cells of the genotypes indicated in the figure legends were plated in 8-well Lab Tek II Chamber 1.5 German coverglass dishes (Thermo Fisher, 155409) in regular growth medium, and incubated for 6 hours. Medium was then changed to Liebowitz-15 without phenol-red (Thermo Fisher, 21083027) supplemented with 10% fetal calf serum, 200 ng/mL doxycycline and 50 nM Sir-DNA (Cytoskeleton, CY-SC007). Cells were incubated for 24 hours prior to imaging. Time lapse images were acquired using Nikon Eclipse Ti2-E inverted microscope (Nikon), equipped with Kinetix sCMOS camera (Photometrics), Spectrax Chroma light engine for fluorescence illumination (Lumencor), or a Nikon Ti / CSU-W1 Spinning Disc Confocal microscope (Nikon), equipped with Photometrics Prime 95B camera (Photometrics) and 3iL35 LaserStack (Intelligent Imaging Innovations Inc). Both systems are equipped with a perfect focus system, and an incubation chamber with 37°C and controlled humidity (OkoLab). Three-dimensional images at multiple stage positions were acquired in steps of 2 μm, every 7 minutes for 10 hours using NIS Elements (Nikon) and 20x Plan Apochromat Lambda objective (NA 0.80, Nikon) or 40x Plan Apochromat Lambda objective (NA 0.95, Nikon). Maximum intensity projections and inverted color profiles of representative examples of mitoses were prepared in Fiji and exported as still images. Analysis of mitotic cells was performed using 3D reconstructions in Fiji. The parameters scored (based on the Sir-DNA signal) were: occurrence of unaligned chromosomes in metaphase, and chromosome segregation errors in anaphase. Analyses of 100 cells per cell line in three biological replicates were documented using Excel and processed and plotted using GraphPad Prism software. Instances where not all the chromosomes were properly aligned on the spindle equator in metaphase and/or anaphase are classified as chromosome alignment errors. Instances where sister chromatids failed to properly separate, either segregating both into the same daughter cell or forming a bridge in anaphase were classified as segregation errors. Numbers reported represent percentage of cells experiencing either abnormality.

### Western blot analysis

For analysis of protein expression levels in HEK T-REx cell lines, cells were typically processed in parallel to cells used for autoregulation assays in 12 or 24 well plates, and protein expression was induced by addition of 1 *μ*g/ml doxycycline for 24–48h. Cells were washed with PBS once and then harvested in PBS, pelleted and lysed in 1% SDS, 100 mM Tris pH8 by boiling for 20 minutes at 95°C. Samples were normalized, separated on 7% or 10% Tris-Tricine based gels, and transferred to 0.2 *μ*m nitrocellulose membrane (BioRad). Membranes were stained with Ponceau S (Sigma), blocked in 5% milk (or 3% BSA for Streptavidin-HRP blots) and incubated with primary antibody at 4°C overnight or for 1h at room temperature as listed below. Signals were detected using HRP-conjugated secondary antibodies and chemiluminescent substrate Pierce ECL or SuperSignal West Pico PLUS (Thermo Fisher). As loading controls, membranes were probed with antibodies against β-actin, RPL8 or GAPDH. Alternatively, the Ponceau S stained membrane is displayed.

For total protein analysis of HeLa cells, parental HeLa T-REx, SCAPER knockout and the indicated rescue cell lines were grown in 6 well plates and treated with 200 ng/ml doxycycline for 24 hours, then washed with PBS and collected by scraping directly in Laemmli buffer. Total cell lysates were boiled for 5 minutes, equal volumes loaded on a Tris-Glycine 4-12% gel (ThermoFisher Scientific, XP04125BOX), and transferred in the presence of 0.1% SDS to nitrocellulose membrane. The membrane was incubated with blocking solution (5% non-fat dry milk in PBS-0.2% Tween 20) and then exposed to primary antibodies against FLAG-tag and GAPDH. The membrane was further incubated with HRP-conjugated secondary antibodies against mouse (ThermoFisher Scientific, 31430) and rabbit (ThermoFisher Scientific, 31460) at 1:10.000 dilution and visualized by ECL (ThermoFisher Scientific, 34580) using an Amersham ImageQuant 800 imaging system.

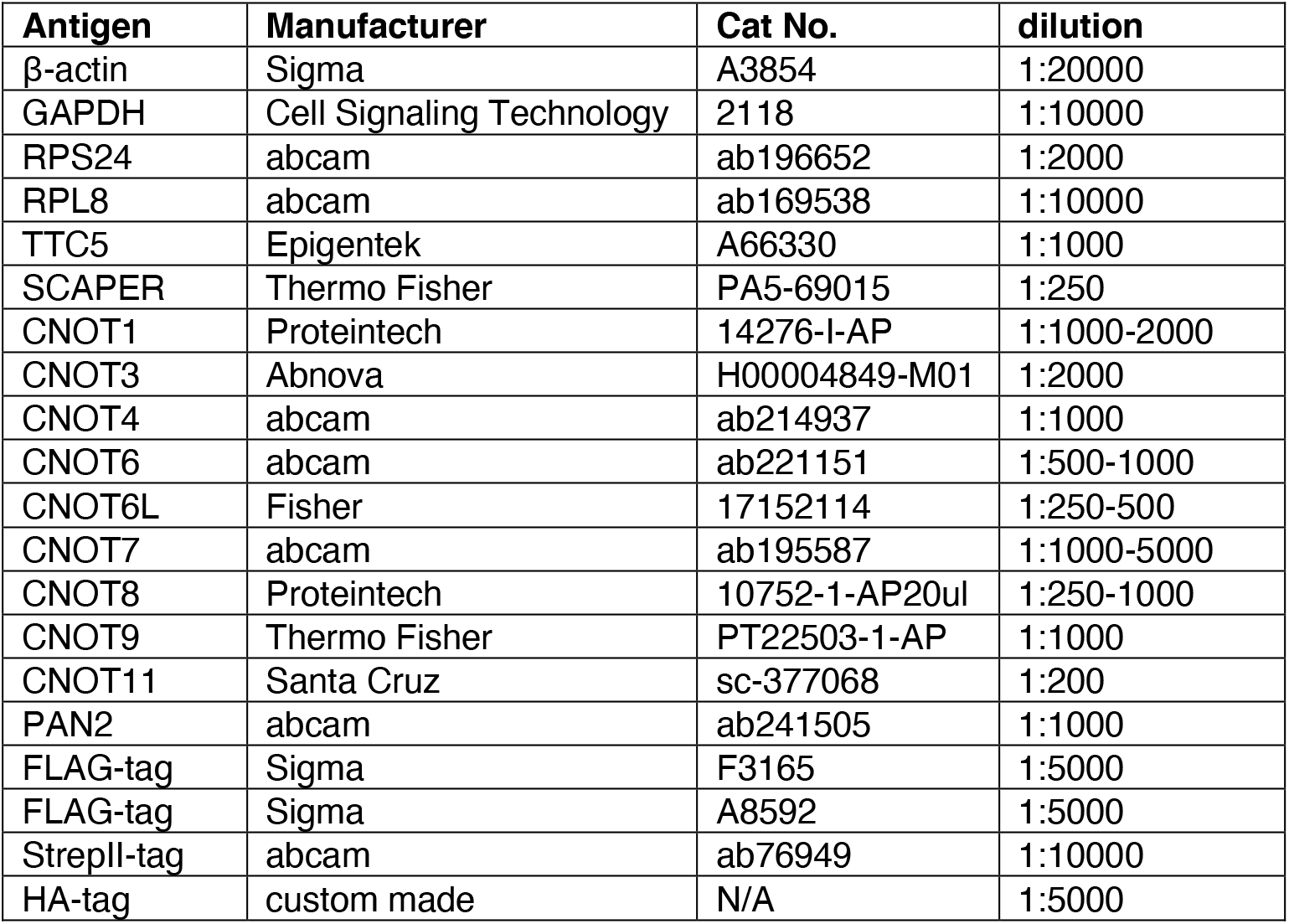
Primary antibodies used in this study.

### Autoregulation assay and RT-qPCR mRNA quantification

For autoregulation assays in HEK T-REx cells, cells were grown to 70–80% confluency (optionally with 1 μg/ml doxycycline for 24–48 h) in 24- or 12-well plates and treated with colchicine (10 μM, Sigma PRH1764), combretastatin A4 (Selleckchem S7783), or as controls (DMSO/regular media) for 3 hours. Cells were washed with PBS, harvested and total RNA was isolated using the RNeasy Plus mini kit (QIAGEN, 74134) as per the manufacturers protocol. 500 ng of total RNA was used to generate cDNA using the iScript cDNA synthesis kit (BioRad 1708891). Samples were diluted ten-fold with nuclease-free water, or kept at higher concentrations to make a standard curve. RT-qPCR was carried out using a ViiA 7 Real-Time PCR System (Thermo Fisher Scientific) and KAPA SYBR Fast qPCR reagents (KAPA Biosystems) as per manufacturer’s instructions. The primer sequences used are listed below. All pairs of primers were annealed at 60°C, and a melt curve performed. PCR products were verified by sequencing. Data was then analyzed using the Quantstudio Real-time PCR software v1.3. Relative standard curve quantification was performed and values were normalized to RPLP1 levels, and to untreated control samples. Experiments include two to three biological replicates, unless stated otherwise. Processing, statistical analysis, and data plotting were performed in Microsoft Excel and GraphPad Prism.

For analysis of previously reported CCR4-NOT substrates, untreated control cDNA samples from siRNA knockdown experiments were reanalysed using TaqMan probes (Thermo Fisher) and TaqMan Fast Advanced Master Mix according to the manufacturer’s protocols. FAM-MGB labelled probes (Cat No 4331182) for TRIB3 (ID: Hs00221754_m1), FZD8 (ID: Hs00259040_s1), LEFTY2 (ID: Hs00745761_s1) and 18S rRNA (ID: Hs99999901_s1) were analyzed in multiplex-reactions with a VIC-MGB labelled GAPDH probe (Cat No 4326317E, ID: Hs99999905_m1). A standard curve was prepared from CNOT1-KD samples. Samples were normalized to GAPDH as an endogenous control for each well, and relative standard curve quantification was performed using the Quantstudio Real-time PCR software v1.3.

For autoregulation assays in HeLa cells, Flp-In TRex HeLa parental, SCAPER knockout and the indicated rescue cell lines were grown to 70–80% confluency in 10 cm dishes and treated with DMSO (control) or combretastatin A4 (100 nM) 4 hours. Cells were harvested and total RNA isolated using the PureLink RNA Mini Kit (Invitrogen, Thermo Fisher, 12183018A) as per manufacturer’s protocol. On column DNase digestion was performed using PureLink DNase Set (Thermo Fisher, 12185010) as per manufacturer’s instructions. 500 ng of total RNA was used to generate cDNA using the SuperScript IV kit (Invitrogen, 18091050) and random hexamer primers following the manufacturer’s protocol. RT-qPCR was carried out using 5 ng of cDNA and 2x PowerUp SYBR Green master mix (Thermo Fisher, A25776) on a thermocycler (BioRad), as per manufacturer’s instructions. Data analysis was performed using the ddCt method^70^. All data were normalized to reference genes RPLP1 or GAPDH, and to DMSO treated controls. Experiments include two biological replicates. Processing and data plotting were performed in R, Microsoft Excel, and GraphPad Prism.

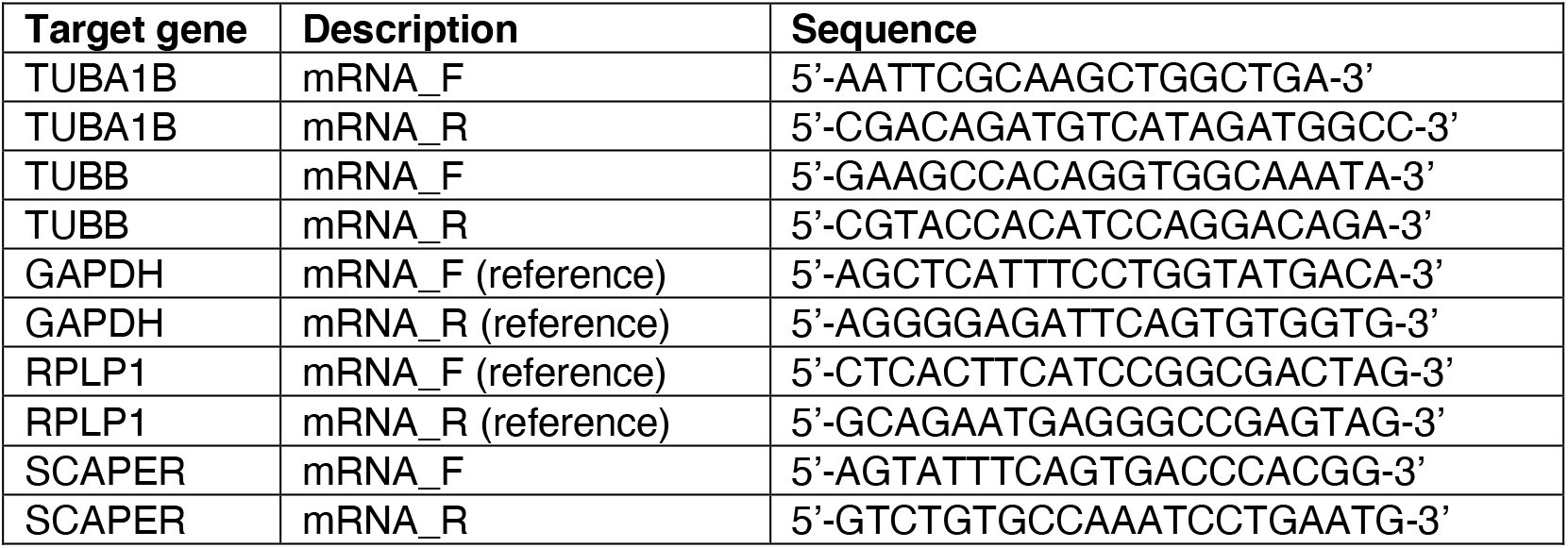
RT-qPCR primers used in this study.

### Pulse labelling of protein synthesis

To measure tubulin autoregulation by pulse labelling of protein synthesis, HEK T-REx wild type or SCAPER KO cells were seeded in 12-well plates and transfected the next day with pcDNA5/FRT/TO rescue plasmids and a puromycin-resistance conferring plasmid (MXS-CMV-PuroR) using TransIT-293 (Mirus). 24 hours after transfection, cells were induced and selected by addition of 1 *μ*g/ml doxycycline and 1 *μ*g/ml puromycin, respectively. 24 hours after induction, cells were treated with 100 nM CA4 (or left untreated) for 3 hours. Cells were then washed with warm PBS and harvested in PBS. 40% of cells were used for total protein analysis, and 60% of cells were resuspended in depletion media lacking FCS and methionine (+/− 100 nM CA4). Cells were starved for 30 minutes at 37°C and pulse labelling was performed for 30 minutes at 37°C by addition of ^35^S-methionine at 100 *μ*Ci/ml. After labelling, cells were pelleted (5000 rpm, 2 min) and lysed in 45 *μ*l digitonin lysis buffer [50 mM HEPES pH7.4, 100 mM KAc, 5 mM MgAc_2_, 1 mM DTT, 1x EDTA-free protease inhibitor cocktail (Roche), 0.01% digitonin] for 10 minutes on ice. Lysates were cleared by centrifugation at maximum speed at 4°C in a table-top centrifuge. 1 *μ*l sample was mixed with sample buffer and separated on 10% Tris-Tricine gels to analyze proteins by autoradiography. Quantification was performed using ImageLab software (BioRad). The tubulin band was normalized to an unrelated band for each lane and then to untreated control samples. Microsoft Excel and GraphPad Prism were used to plot data. Two independent replicates were averaged.

### Biotin proximity labelling procedure

For biotin proximity labelling experiments^35,71^, TurboID-FLAG was fused to the N-terminus of TTC5 or SCAPER (WT or mutants) and cloned into pcDNA5/FRT/TO vectors. TTC5 or SCAPER KO HEK T-REx cell lines were rescued by stable integration of TurboID constructs, which were functional in autoregulation assays. To avoid strong overexpression, leaky expression from the doxycycline-inducible promoter was used for TurboID-TTC5 expression, and TurboID-SCAPER was induced with 2 ng/ml doxycycline for 48 hours. Parental cell lines without TurboID constructs served as specificity controls for mass spectrometry.

To isolate biotinylated proteins for mass spectrometry analysis, cells were seeded in 150 mm plates and grown to ~ 80% confluency. For TurboID-TTC5, two plates per replicate were pretreated with DMSO (control), colchicine (10 μM, Sigma PRH1764) or nocodazole (10 μM, Sigma SML1665) for 30 minutes and biotin (APExBIO A8010) was added at 50 μM and incubated for another 2.5 hours. For SCAPER, one plate of cells per replicate was treated with DMSO (control) or combretastatin A4 (Selleckchem S7783) for 30 minutes and biotin was added at 50 μM and incubated for another 30 minutes. Cells were washed once in ice-cold PBS, pelleted, and cytosolic extracts were prepared by lysis in 1 ml digitonin lysis buffer per 150 mm plate for 10–15 min on ice [50 mM HEPES pH7.4, 100 mM Kac (400 mM KAc for TurboID-SCAPER samples), 5 mM MgAc_2_, 1 mM DTT, 1x EDTA-free protease inhibitor cocktail (Roche), 0.01% digitonin]. Lysates were cleared by centrifugation at maximum speed at 4°C in a table-top centrifuge. Lysates were then incubated on a rotating wheel with ~ 50 μl of streptavidin-coupled magnetic beads (Pierce 88817) for 2 hours at 4°C. Beads were then washed with 1 ml each of physiological salt buffer [PSB: 50 mM HEPES pH7.4, 100 mM KAc (400 mM for TuroboID-SCAPER samples), 2 mM MgAc_2_] with 0.01% digitonin, wash buffer 1 (1% SDS, 10 mM Tris-HCl pH8), wash buffer 2 (1 M NaCl, 10 mM HEPES pH7.4, 0.01% digitonin), and wash buffer 3 (2 M urea, 10 mM Tris-HCl pH8, 0.01% digitonin). To remove detergent, beads were washed twice with 100 μl 50 mM Tris-HCl pH8, 150 mM NaCl and transferred to a new tube with the last step. Beads were then stored in 20 μl 50 mM Tris-HCl pH8, 150 mM NaCl for mass spectrometry analysis, or eluted with 20 μl sample buffer supplemented with 2 mM biotin for 5 minutes at 95°C for analysis by SDS-PAGE. For mass spectrometry analysis, two or three biological replicates were processed for each condition.

For western blot validation of SCAPER biotinylation by TurboID-TTC5, expression in the indicated cell lines was induced with 1 μg/ml doxycycline and cells were transfected with a pcDNA3-SCAPER-FLAG construct using TransIT293 (Mirus) in 10 cm dishes. All plates were treated with colchicine (10 μM 2.5 hours) and biotin was added for another 30 minutes (50 μM). Biotinylated proteins were isolated as described above.

### Quantitative proteomics procedures

#### On-bead digestion

Proteins bound to beads were reduced with 2 mM DTT in 2 M urea buffer and sequencing grade trypsin (Promega) was added to a final concentration of 5 ng/*μ*l. After incubation for 3 h at 25°C, supernatants were transferred to fresh eppendorf tubes. Beads were washed once with 2M urea buffer, once with 1M urea buffer, and the washes were combined with the corresponding supernatants. Samples were then alkylated with 4 mM iodoacetamide (IAA) in the dark at 25°C for 30 min. An additional 0.1 *μ*g of trypsin (Promega) was added to the samples and digested over night at 25°C. Samples were acidified to 0.5% formic acid (FA) and desalted using home-made C18 (3M Empore) stage tips filled with 4 *μ*l of Poros Oligo R3 resin (Thermo Fisher). Bound peptides were eluted sequentially with 30%, 50% and 80% acetonitrile (MeCN) in 0.5% FA and lyophilized.

#### Tandem mass tag (TMT) labeling

Dried peptides from each condition were resuspended in 15 *μ*l of 200 mM HEPES, pH 8.5. 7.5 *μ*l of TMTpro 18-plex reagent (Thermo Fisher Scientific), reconstituted in anhydrous acetonitrile according to manufacturer’s instructions, was added and incubated at room temperature for 1 h. The labeling reactions were terminated by incubation with 1.5 *μ*l of 5% hydroxylamine for 30 min. Labeled samples for each condition were pooled into one sample, and MeCN was removed by vacuum centrifugation. TMT-labeled peptides were desalted and then fractionated with home-made C18 stage tip using 10 mM ammonium bicarbonate and increasing acetonitrile concentration. Eluted fractions were acidified, partially dried down in a speed vac and used for LC-MS/MS.

#### Mass spectrometry analysis

The fractionated peptides were analysed by LC-MS/MS using a fully automated Ultimate 3000 RSLC nano System (Thermo Fisher Scientific) fitted with a 100 *μ*m × 2 cm PepMap100 C18 nano trap column and a 75 *μ*m × 25cm, nanoEase M/Z HSS C18 T3 column (Waters). Peptides were separated using a binary gradient consisting of buffer A (2% MeCN, 0.1% FA) and buffer B (80% MeCN, 0.1% FA). Eluted peptides were introduced directly via a nanospray ion source into a Q Exactive Plus hybrid quardrupole-Orbitrap mass spectrometer (MS2, TurboID-TTC5 samples) or Orbitrap Eclipse mass spectrometer (RTS-MS3, TurboID-SCAPER samples), both from Thermo Fisher Scientific. The Q Exactive Plus mass spectrometer was operated in standard data dependent mode, performed MS1 full-scan at m/z = 380-1600 with a resolution of 70K, followed by MS2 acquisitions of the 15 most intense ions with a resolution of 35K and NCE of 29%. MS1 target values of 3e6 and MS2 target values of 1e5 were used. Dynamic exclusion was enabled for 40s.

For the RTS-MS3 experiment (TurboID-SCAPER samples), MS1 spectra were acquired using the following settings: Resolution=120K; mass range=400-1400m/z; AGC target=4e5 and dynamic exclusion was set at 60s. MS2 analysis were carried out with HCD activation, ion trap detection, AGC=1e4; NCE=33% and isolation window =0.7m/z. RTS of MS2 spectrum was set up to search uniport Human proteome (2021), with fixed modifications cysteine carbamidomethylation and TMTpro 16plex at N-terminal and lysine residues. Met-oxidation was set as variable modification. Missed cleavage=1 and maximum variable modifications=2. In MS3 scans, the selected precursors were fragmented by HCD and analyzed using the orbitrap with these settings: Isolation window=1.3 m/z; NCE=55; orbitrap resolution=50K; scan range=110-500 m/z and AGC=1e5.

#### Data analysis

The acquired LC-MS/MS raw files, were processed using MaxQuant^72^ with the integrated Andromeda search engine (v1.6.6.0 or v1.6.17.0). MS/MS spectra were quantified with reporter ion MS2 or MS3, and searched against Human Reviewed UniProt Fasta database (downloaded in 2019). Carbamidomethylation of cysteines was set as fixed modification, while methionine oxidation and N-terminal acetylation (protein) were set as variable modifications. Protein quantification requirements were set at 1 unique and razor peptide. In the identification tab, second peptides and match between runs were not selected. Other parameters in MaxQuant were set to default values.

The MaxQuant output file (proteinGroups.txt) was then processed with Perseus software (v1.6.6.0 or v1.6.15.0). After uploading the matrix, the data was filtered to remove identifications from reverse database, identifications with modified peptide only, and common contaminants. Data were log2-transformed, a valid value filter was applied and missing values for remaining proteins were imputed with standard settings. Data were then exported for further processing in MS Excel, where intensity values were normalized to bait protein levels for each sample, except for untagged control samples. Background binders were filtered if intensities were less than 4-fold enriched in any sample over an untagged cell line (TurboID-TTC5), or if average intensity in TurboID samples was less than 2-fold enriched over untagged control levels (TurboID-SCAPER). A two-tailed t-test was used to calculate p-values between sample groups. For TurboID-TTC5 versus K97A comparison, values from all conditions (DMSO, colchicine, nocodazole, two replicates each) were used for statistics, because SCAPER binding was independent of treatments. For TurboID-SCAPER, three replicates each of TurboID-SCAPER +/− CA4, and two replicates for E620Δ + CA4 samples were analyzed. Data were plotted in GraphPad Prism.

### Recombinant protein purification

WT and mutant 6xHis-TEV-Twin-Strep-tagged TTC5 (“Strep-TTC5”) were purified from E. coli cells as described^17^. Briefly, BL21 DE3 cells were transformed with the respective pET28a plasmids and grown at 37°C in LB containing 50 *μ*g/ml kanamycin. Induction was with 0.2 mM IPTG at an A600 of 0.6 at 16°C overnight. Bacterial lysate was prepared by sonication (Sonics Vibracell) in 25 ml cold lysis buffer [500 mM NaCl, 20 mM imidazole, 1 mM TCEP, 1x EDTA-free protease inhibitor cocktail (Roche), and 50 mM HEPES, pH7.4] per litre of cells. Clarified bacterial lysates from a 1 l culture were bound to a 0.5 ml column of Ni-NTA resin (Qiagen) by gravity flow. Columns were washed with ~40 column volumes of lysis buffer and eluted with 250 mM imidazole in lysis buffer. The eluate was then bound to a 200 *μ*l column of Streptactin Sepharose (IBA 2-1201-010). After extensive washing with 500 mM NaCl, 1 mM TECP and 50 mM HEPES, pH 7.4, TTC5 protein was eluted with 400 *μ*l washing buffer containing 50 mM biotin and dialyzed against dialysis buffer (500 mM NaCl, 25 mM HEPES, pH 7.4).

Recombinant N- or C-terminally FLAG-tagged SCAPER was purified from Expi293 cells. Briefly, 100 ml cells were transfected with pcDNA3 or pcDNA5 plasmids encoding SCAPER constructs using polyethyleneimine-Max (made in-house) and grown for 72 hours for protein expression. Cells were pelleted and lysed in 10 ml lysis buffer [50 mM HEPES pH7.4, 400 mM KAc, 2 mM MgAc_2_, 0.01% digitonin, 1 mM DTT, 1x EDTA-free protease inhibitor cocktail (Roche)] using a dounce homogenizer. Cleared lysates were incubated with 250 *μ*l anti-FLAG resin (Sigma) with rotation for 2 hours at 4°C. The resin was then transferred to a gravity flow column (BioRad) and washed with 80 column volumes of lysis buffer, and 20 column volumes of wash buffer (50 mM HEPES pH7.4, 400 mM KAc, 2 mM MgAc_2_). Proteins were eluted in two column volumes of 0.2 mg/ml 3xFLAG peptide (Sigma) and dialyzed against 50 mM HEPES pH 7.4, 400 mM KAc, 2 mM MgAc_2_.

### Pull-down assays

For pull-downs of recombinant SCAPER by TTC5, proteins were mixed at 100 nM (SCAPER) or 150 nM (TTC5) final concentration in 400 *μ*l reactions in IP buffer (50 mM HEPES pH7.4, 100 mM KAc, 5 mM MgAc_2_, 1 mM DTT, 0.01% digitonin). Reactions were incubated rotating for 1 hour at 4°C, 5 μl of streptactin magnetic agarose beads (IBA 2-4090-010) were added and samples were incubated another 1 hour. Beads were washed five times with 400 μl IP buffer and transferred to a new tube with the last step. Proteins were eluted with sample buffer, separated by SDS-PAGE, and gels were stained with Coomassie brilliant blue.

### In vitro transcription and translation

All in vitro transcription of tubulin constructs utilized PCR product as template and were carried out as described^17^. The 5’ primer contained the SP6 promoter sequence and anneals to the CMV promoter of pCDNA3.1. The 3’ primers anneal at codon 54-60 or 84-90 of tubulin and contain extra sequence encoding MKLV to generate 64-mer or 94-mer nascent chains, respectively. Transcription reactions were carried out with SP6 polymerase (NEB) for 1 hour at 37°C. Transcription reactions were directly used for in vitro translation in a homemade rabbit reticulocyte lysate (RRL)-based translation system as previously described^73,74^, optionally in the presence of ^35^S-methionine. Recombinant Strep-TTC5 (100–250 nM) or FLAG-SCAPER (100–250 nM) proteins were included in the translation reactions as indicated. Translation reactions were at 32°C for 15 minutes, or 30 minutes for large-scale reactions for structural analysis. For analysis of total translation level of nascent chains, a 1 μl aliquot of the translation reaction was mixed with protein sample buffer and analyzed by SDS-PAGE gel electrophoresis and autoradiography.

For analysis of ribosome nascent chains (RNCs), translation products were pulled down via the Twin-Strep-tag on TTC5 using Streptactin Sepharose (IBA 2-1201-010) for 2 hours at 4°C. Beads were washed four times with PSB (50 mM HEPES pH7.4, 100 mM KAc, 2 mM MgAc_2_) and eluted with 50 mM biotin in PSB for 30 minutes on ice. Elutions were analyzed by SDS-PAGE followed by SYPRO Ruby staining (Thermo Fisher), western blotting, or autoradiography. Alternatively, translation reactions were separated on linear 10–50% sucrose gradients (55,000 rpm, 20 min) and analyzed as above.

### Structural analysis of TTC5-SCAPER-ribosome complexes

#### Cryo-EM grid preparation and data collection

Affinity purified ribosomes at a concentration of ~ 65 nM (A_260_ of 3.2) were vitrified on UltrAuFoil R1.2/1.3 300-mesh grids (Quantifoil) coated with graphene oxide (GO). For GO coating, gold grids were washed with deionized water, dried and subsequently glow-discharged for 5 min with an Edwards glowdischarger at 0.1 torr and 30 mA. 3 *μ*l of a 0.2 mg/ml GO suspension in deionized water (Sigma) was pipetted onto the glow-discharged grids and incubated for 1 min. Next the GO solution was blotted away and the grids were washed 3x by dipping into 20 *μ*l deionized water drops followed by blotting (washed twice the top-side and once the bottom-side). 3 *μ*l sample was pipetted onto the grids, blotted for 5.5 sec, −15 blot force, 0 sec wait at 100% humidity, 4°C, Whatman 595 blotting paper, with a Vitrobot Mark IV and plunge frozen into liquid ethane. Grids were stored in liquid nitrogen until data-collection. The dataset was collected with a Gatan K3 camera on Titan Krios4 microscopes at eBIC (Diamond) in super-resolution counting mode and binning 2, using EPU software in faster acquisition mode (AFIS) yielding 20932 micrographs (105000x magnification, pixel size= 0.829 Å, total dose 44.7 e^-^/Å^2^, 44 frames, resulting in 1 e^-^/Å^2^ dose/frame). Refer to Supplementary Table S1 for data collection statistics.

#### Cryo-EM data processing

Datasets were processed with RELION 4. Raw movies were corrected with MotionCor (5×5 patches), followed by CTF correction using CTFFIND-4.1. Particles were picked using low-pass filtered 80S ribosomes as a 3D reference, resulting in 1227269 initial particles, which were used for initial 2D classification. Good 2D classes (696074 particles) were selected and subjected to 3D classification without alignment using data to 8.29 Å, which resulted in 559080 high-resolution 80S particles. Particles were then re-extracted at 1.32 Å/pixel and 3D refined yielding an overall resolution of 2.89 Å. To select TTC5 and SCAPER bound 80S, we performed a focused classification with signal subtraction (FCwSS) around TTC5 and SCAPER without alignment, resulting in 22610 particles. These particles were then extracted at full pixel size (0.829 Å) and we performed Bayesian polishing and CTF refinement [(anisotropic) magnification estimation followed by CTF parameter fitting (fit defocus, astigmatism and B-factor per particle)], resulting in a map with an overall resolution of 2.89 Å. These particles were subjected to three different FCwSS. First, we performed a FCwSS around the expansion segment contacting SCAPER, yielding two classes with density corresponding to the expansion segment, resulting in two maps with an overall resolution of 3.10 Å (9158 particles) and 3.17 Å (7370 particles), respectively. Second, we performed a FCwSS around the P-site tRNA, resulting in a map with 3.24 Å resolution (5424 particles). Third, we did a FCwSS around SCAPER to remove some non-SCAPER containing particles, which resulted in a map with an overall resolution of 2.95 Å (18949 particles). The 40S subunit was in several rotation states, so we subtracted the 40S and focused on the 60S. This step resulted in an overall resolution of 2.84 Å after 3D refinement for the 60S subunit bound by TTC5 and SCAPER.

#### Model building, refinement and validation

The molecular model from PDB 6T59 (60S bound to TTC5 and tubulin nascent chain) was split into two groups, which were individually docked into the 2.8 Å post-processed map using UCSF Chimera (version 1.15)^75^. TTC5 and the nascent chain from PDB 6T59 were deleted and replaced by an Alphafold2 model^37,38,76^ of TTC5 bound to the β-tubulin nascent chain. The SCAPER model was also derived from Alphafold2. Similar to the two groups of PDB 6T59, TTC5-β-tubulin nascent chain and SCAPER were individually docked in Chimera. Subsequently, all chains were manually adjusted into the original, or suitably blurred maps (B factors of 60 to 100) using Coot (version 0.9.6, Marina Bay)^77^. TTC5–β-tubulin nascent chain and SCAPER were merged into group 2 of PDB 6T59. In Phenix (version 1.20-4459-000)^78^, the 2 groups were first combined using *iotbx.pdb.join_fragment_files* and then *phenix.real_space_refine* was used to perform real space refinement of the resulting model with default settings and the following additions: *phenix.elbow* was used to automatically obtain restraints for all non-standard RNA bases and ligands; nonbonded weight of 1000 was used; rotamer outliers were fixed using the Fit option ‘outliers_or_poormap’ and the Target was set to ‘fix_outliers’; and finally 112 processors were used to speed up the calculations. Refer to Supplementary Table S1 for processing, refinement and model statistics.

#### Molecular graphics

Map and model figures were generated using UCSF Chimera (version 1.15)^75^, UCSF Chimera X (version 1.3)^79^ and PyMOL (Molecular Graphics System, version 2.4, Schrödinger, LLC). 2D class averages were generated in RELION 4.0 and the FSC curve was plotted using GraphPad Prism.

## Structural modelling

Structure predictions were performed with AlphaFold2 through a local installation of Colabfold 1.2.0^80^, using MMseqs2^81^ for homology searches and AlphaFold2^37^ or AlphaFold2 multimer^38^ for the predictions of single or multiple chains, respectively.

## Data availability

The cryo-EM map will be deposited to the EMDB and atomic coordinates will be deposited to the Protein Data Bank. Mass spectrometry raw data will be deposited to the Proteomics Identifications Database. All other data are available in the manuscript or the supplementary materials.

## Supplemental information

Supplementary Figures S1–S9.

Supplementary Tables S1–S3.

---

**Fig. S1.**
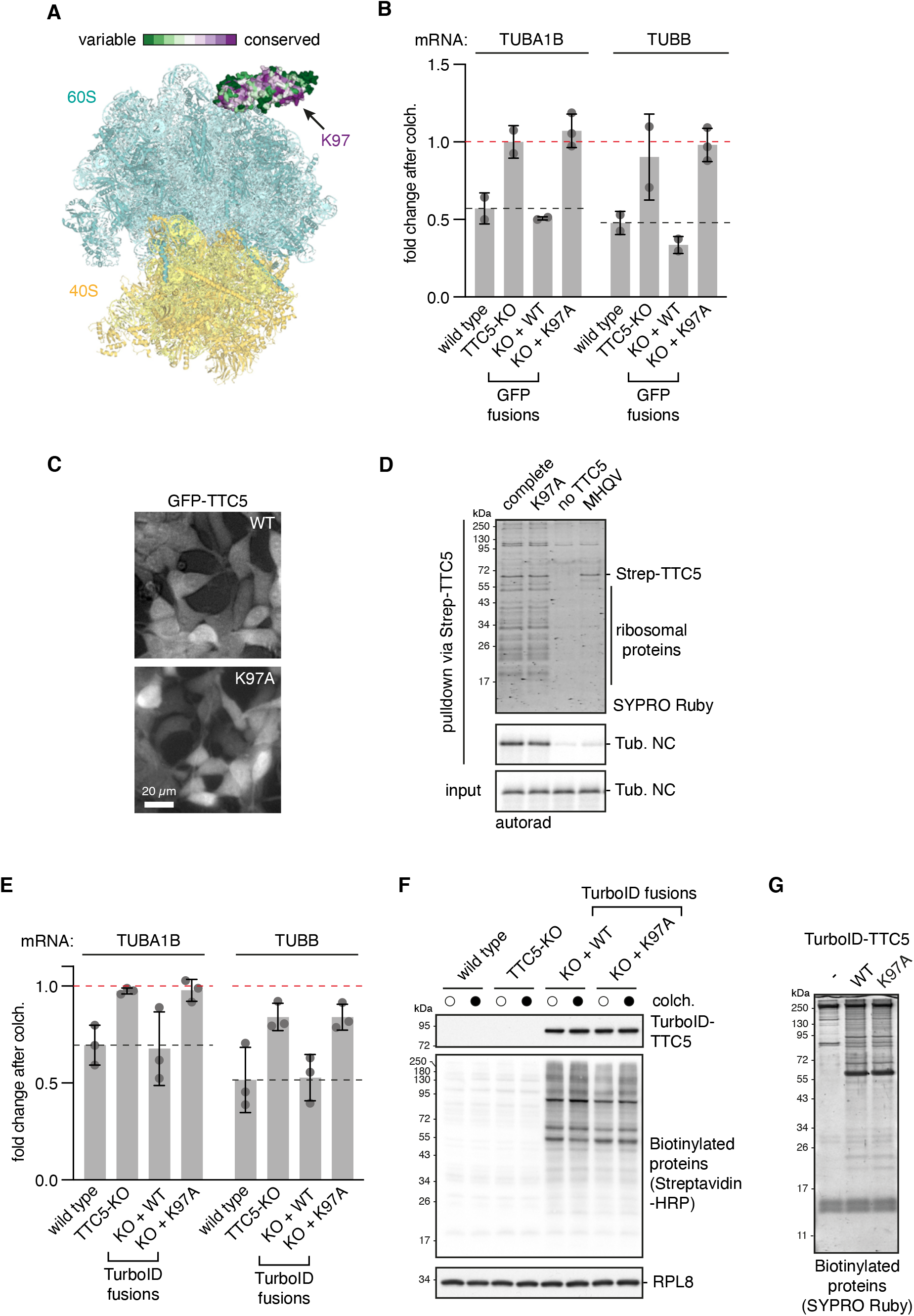
The TTC5 K97A mutation abolishes tubulin autoregulation, related to Fig. 1. (A) Model of TTC5 bound to the ribosome containing a 64 amino acid β-tubulin nascent chain from a previously published structure^17^ (PDB: 6T59) with the 40S subunit docked for reference. ConSurf^82^ was used to display the residue conservation of TTC5 on the surface. The highly conserved surface patch containing the K97 residue is indicated. This patch is not involved in either ribosome binding or nascent tubulin recognition, leading us to a hypothesis in which it is a docking site for downstream factors. (B) Tubulin autoregulation assay as described in Fig. 1B for TUBB, with TUBA1B reproduced from Fig. 1B for comparison. The red dashed line indicates the starting tubulin mRNA level prior to colchicine, arbitrarily set to a value of 1. The black dashed lines indicate the fold change in wild type (WT) cells for each tubulin subunit. Both TUBA1B and TUBB mRNA were quantified in all experiments in the manuscript. Because the results were comparable for both mRNAs throughout, we display only one tubulin gene in most figures for brevity and clarity. TTC5-KO cells were complemented by stable inducible expression with either GFP-tagged wild type (WT) TTC5 or the K97A mutant. (C) The complemented GFP-tagged cell lines from panel B were imaged using a wide-field microscope (Life Technologies EVOS FL) to verify that expression and localization were similar for both constructs. (D) Reconstitution of TTC5 recruitment to tubulin ribosome nascent chains (RNCs). 94-residue β-tubulin (TUBB) nascent chains were produced in rabbit reticulocyte lysates in the presence of recombinant WT or K97A mutant Twin-Strep-tagged TTC5 (Strep-TTC5) as indicated. “MHQV” indicates a β-tubulin construct in which its TTC5-interacting MREI motif has been mutated. TTC5-associated proteins were enriched via its Strep tag, separated by SDS-PAGE and visualized by SYPRO Ruby staining for total protein (top) or autoradiography for the β-tubulin nascent chain (Tub. NC, bottom). (E) Autoregulation assay with the indicated WT, TTC5-KO, or TurboID-TTC5 rescue cell lines. (F) Western blot analysis of cell lines used in panel A. Cells were pre-treated with 10 *μ*M colchicine (colch.) for 30 minutes as indicated by filled circles, and 50 μM biotin was added for 2.5 h (all samples). (G) Biotinylated proteins enriched from parental, TurboID-TTC5 WT or K97A cell lines as detected by SYPRO Ruby staining. Samples prepared in the same way were used for the mass spectrometry experiment in Fig. 1C. 50 μM biotin was added 2.5h prior to harvesting of cells. To avoid over-expression of TurboID-TTC5 constructs in biotinylation experiments, we exploited the basal leaky expression from the tetracycline-regulated CMV promoter seen in the absence of added doxycycline. Bar graphs show mean +/− SD.

**Figure S2.**
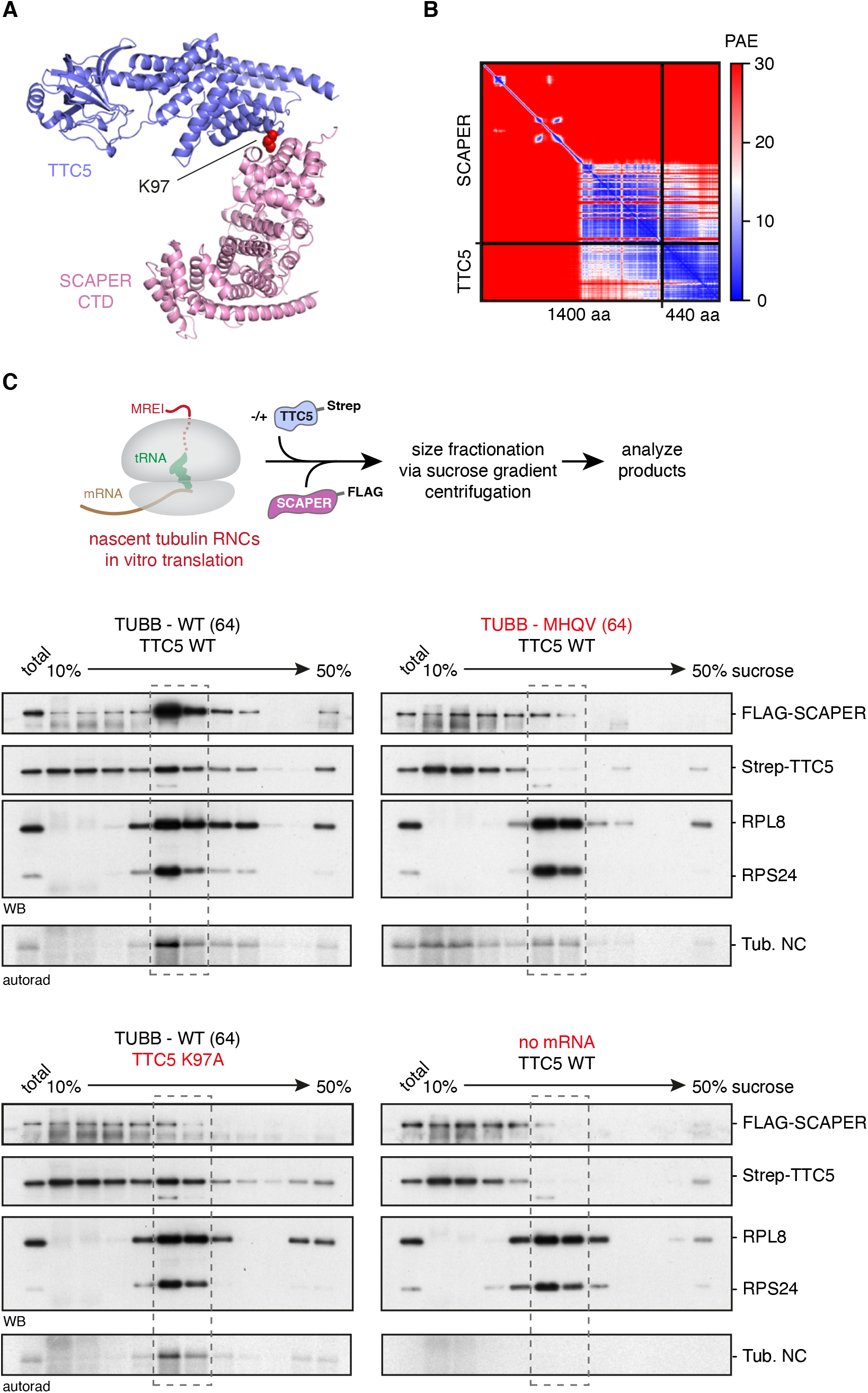
TTC5 recruits SCAPER to tubulin ribosome nascent chain complexes, related to Fig. 2. (A) Top-ranked Alphafold2 (AF2) multimer prediction of the interaction between TTC5 and SCAPER. The SCAPER C-terminal domain (CTD, residues 857–1400) is shown, with the rest of the molecule omitted for clarity. The TTC5 K97 residue is highlighted in red spheres. (B) Predicted Aligned Error (PAE) matrix of the TTC5-SCAPER AF2 multimer prediction shown in panel B. Lower PAE values (blue) indicate higher confidence in the relative positions and orientation between residue pairs in the model. Note that a strong signal is observed between TTC5 and the CTD of SCAPER. (C) 64-residue β-tubulin (TUBB) nascent chain complexes were produced by in vitro translation in the presence of SCAPER and TTC5 as in Fig. 2C. Reactions were then centrifuged through 10–50% sucrose gradients and fractions were analysed by SDS-PAGE, western blotting (WB), and autoradiography (autorad). An aliquot of the total reaction is analyzed in the first lane of each gradient. The peak ribosome-containing fractions, predominantly monosomes, are indicated by dashed boxes. Both the WT β-tubulin sequence and WT TTC5 are required for efficient recruitment of SCAPER to RNCs.

**Figure S3.**
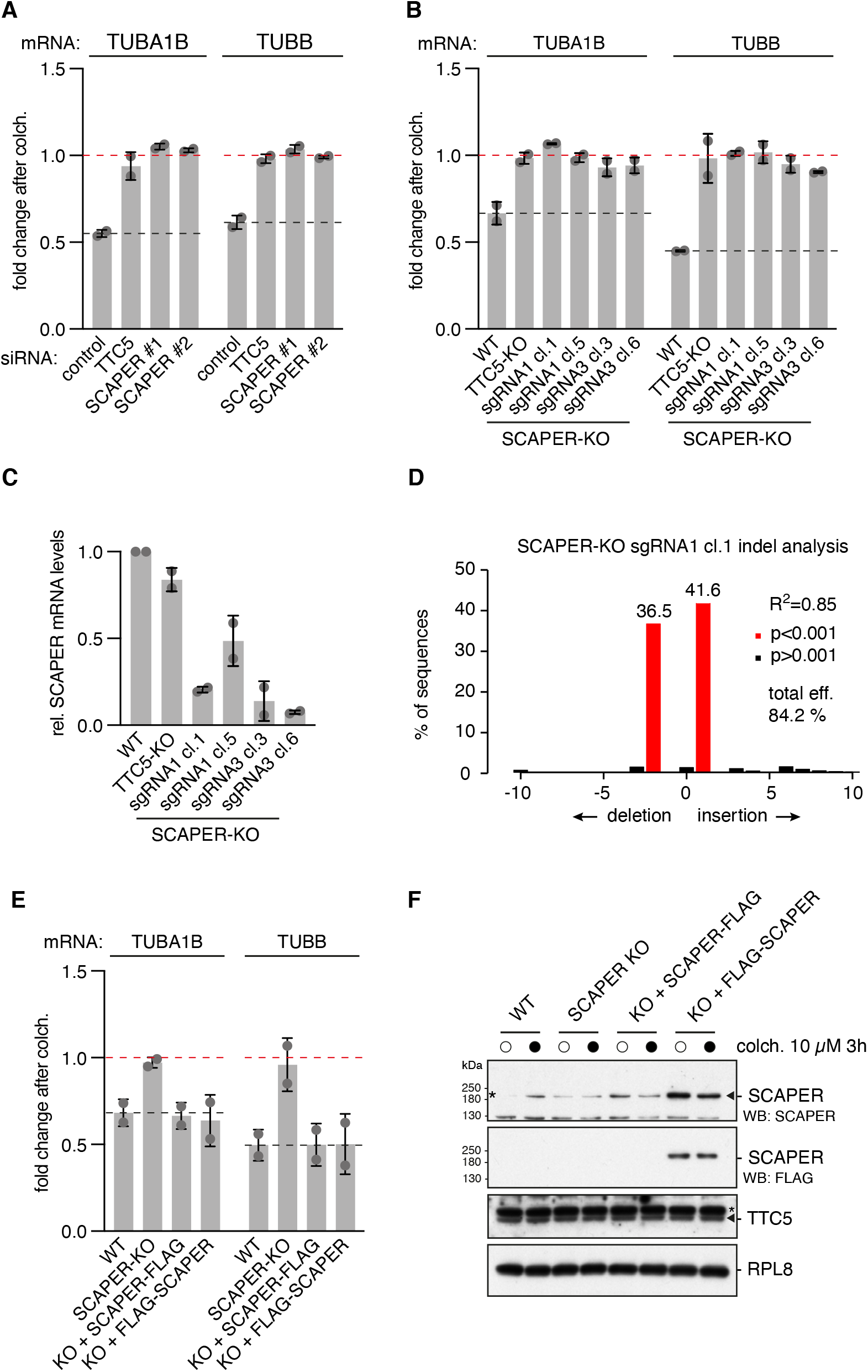
SCAPER is required for tubulin autoregulation, related to Fig. 2. (A) Tubulin mRNA levels were determined in the qPCR-based autoregulation assay for cells transfected with the indicated siRNAs for 72h. Samples from colchicine-treated cells (10 *μ*M for 3h) were normalized to untreated control samples, arbitrarily set to a value of 1 (red dashed line). The black dashed lines indicate the fold change in wild type (WT) cells for each set of samples. (B) Autoregulation assays were performed using the indicated cell lines or SCAPER KO clones. (C) SCAPER mRNA levels were quantified using RT-qPCR in the indicated cell lines and SCAPER KO clones. Reduced mRNA levels suggest that premature termination codons resulting from indels render SCAPER mRNA subject to nonsense-mediated decay. (D) Indel analysis for SCAPER KO sgRNA1 cl.1 using genomic DNA sequencing followed by TIDE analysis^69^. The same clone was used throughout the rest of the study. (E) Autoregulation assay demonstrating that both C- or N-terminally FLAG-tagged SCAPER constructs rescue the autoregulation defect of SCAPER-KO cells. (F) Cell lines used in panel E were subjected to total protein analysis using western blotting. Note that the α-SCAPER antibody recognizes SCAPER only when over-expressed (triangle), but also crossreacts with a co-migrating background band (asterisk) that appears to be stronger than the endogenous SCAPER signal. The C-terminally tagged SCAPER-FLAG constructs, although expressed sufficiently to rescue autoregulation (panel E), were not readily detected on western blots. For this reason, N-terminally tagged FLAG-SCAPER constructs were used for all subsequent experiments. TTC5 (triangle) was unchanged in SCAPER KO cells. Bar graphs show mean +/− SD, except for panel D.

**Figure S4.**
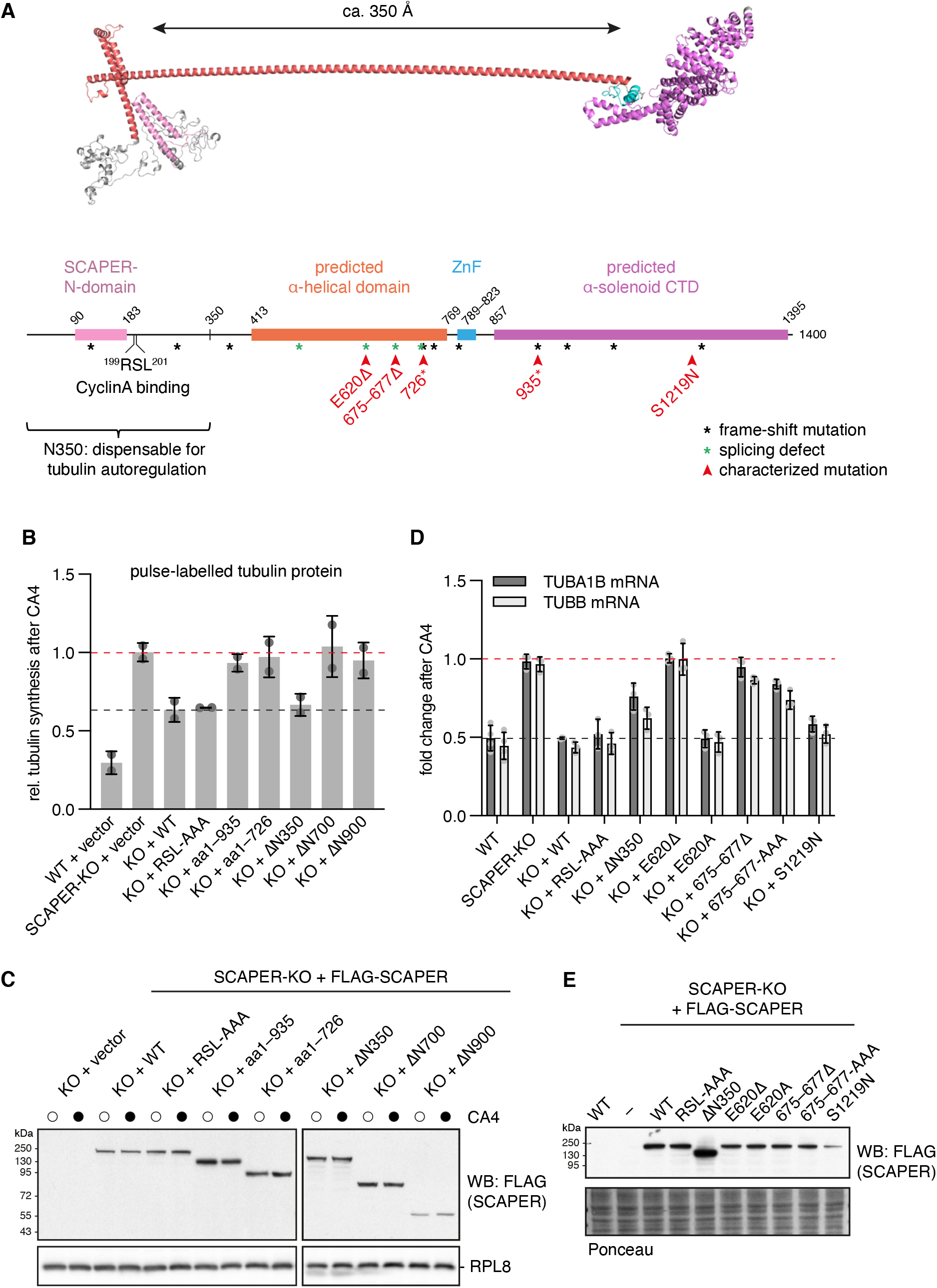
The SCAPER α-helical and C-terminal domains are critical for tubulin autoregulation, related to Fig. 2. (A) Top: Structural model of SCAPER predicted by trRosetta^83^. The displayed model was stitched from two overlapping predictions. Subdomains are color-coded as in the schematic below. Bottom: Schematic of the SCAPER domain architecture with annotated and predicted features indicated (drawn to scale). Pathologic mutations^40–42^ characterized in autoregulation assays in this study are indicated by red arrow heads. Positions of pathologic frameshift and splice site mutations are indicated by black and green asterisks, respectively. RSL: cyclin A binding motif (Arg^199^-Ser^200^-Leu^201^); ZnF: Zinc finger; CTD: Carboxy-terminal domain. (B) Tubulin autoregulation was tested by pulse-labelling of newly synthesized proteins with ^35^S-methionine. Indicated cell lines were transiently transfected with FLAG-SCAPER encoding plasmids. Plotted is the ratio of ^35^S-labelled tubulins from cells after 3h combretastatin A4 (CA4) treatment versus untreated control conditions. RSL-AAA: mutation of the cyclin A binding site (Arg^199^-Ser^200^-Leu^201^) to alanines. Numbers refer to SCAPER amino acid positions, ΔN denotes deletion of the N-terminus up to the indicated residue. Mean +/− standard deviation from two independent experiments. We note that pathologic alleles with truncations after residue 726 and 935 have been reported to cause disease phenotypes such as intellectual disability, a Bardet-Biedl syndrome-like illness, and male infertility^41,43^. (C) Total protein analysis by western blotting of cells used in panel B. (D) Autoregulation assay with WT, SCAPER-KO and the indicated stable rescue cell lines. Note that part of the data is reproduced from Fig. 2D for comparison. Mean +/− SD from at least 3 replicates is plotted. (E) Western blot analysis of SCAPER expression levels of cell lines used in panel D and Fig. 2D. Note that the SCAPER-S1219N variant shows lower expression levels. This indicates that the mutation might destabilize the protein, a potential explanation for the pathologic phenotype it has in a heterozygous patient with the E620Δ mutation on the second allele^40^. Under overexpression conditions, SCAPER-S1219N is functional for autoregulation (see panel D).

**Figure S5.**
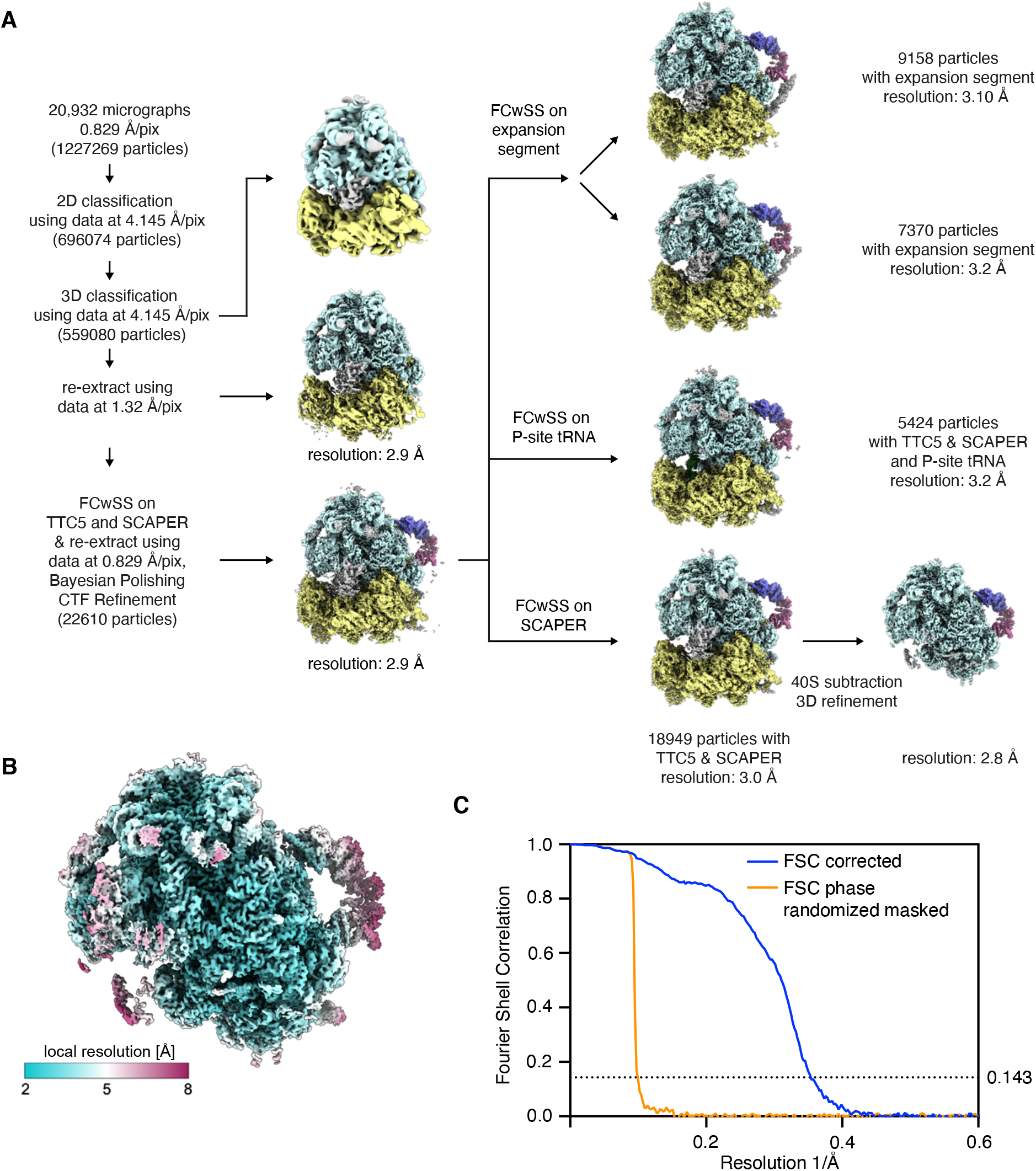
Cryo-EM analysis of SCAPER bound to TTC5 and 80S ribosomes, related to Fig. 3. (A) RELION 4.0 classification and processing workflow for reconstruction of ribosomal particles from cryo-EM micrographs. FCwSS: Focussed classification with signal subtraction. (B) Map coloured according to the local resolution after final FCwSS around SCAPER and subtraction of the 40S subunit. (C) Gold-standard Fourier shell correlation (FSC) curve (blue) of the final 60S-TTC5-SCAPER map illustrating an overall resolution of 2.8 Å. The phase-randomized, masked FSC curve (orange) is also shown.

**Figure S6.**
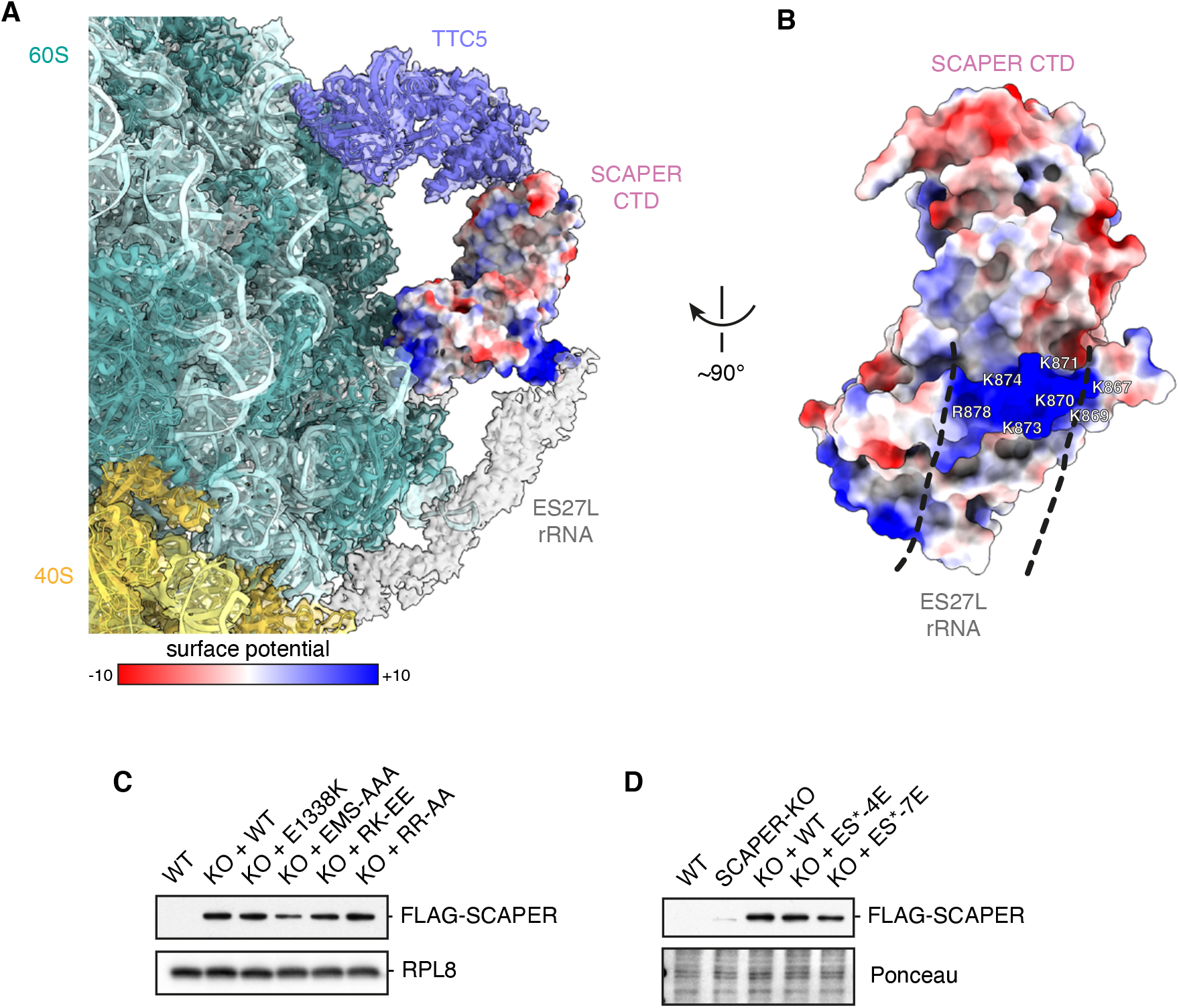
SCAPER contacts with 28S rRNA expansion segment ES27L, related to Fig. 3. (A) View of the cryo-EM map with electrostatic surface potential [in kcal/(mol\**e*)] displayed on the SCAPER CTD model. Note the two highly basic surface patches on SCAPER that contact the 60S body and ES27L rRNA. (B) Side-view of the SCAPER-CTD model from panel A with the path of the density for the rRNA expansion segment ES27L marked by dashed lines. Positively charged residues that were targeted by mutagenesis are indicated. (C, D) Western blot analysis of expression levels of FLAG-SCAPER constructs used in Fig. 3E and 3F, respectively. EMS-AAA: E1338A, M1339A, S1340A; RK-EE: R907E, K910E; RR-AA: R934A, R941A. SCAPER mutations targeting expansion segment contacts were as follows: ES*-4E: K867E, K870E, K873E and K874E; ES*-7E: as ES*-4E plus K869E, K871E and R878E.

**Figure S7.**
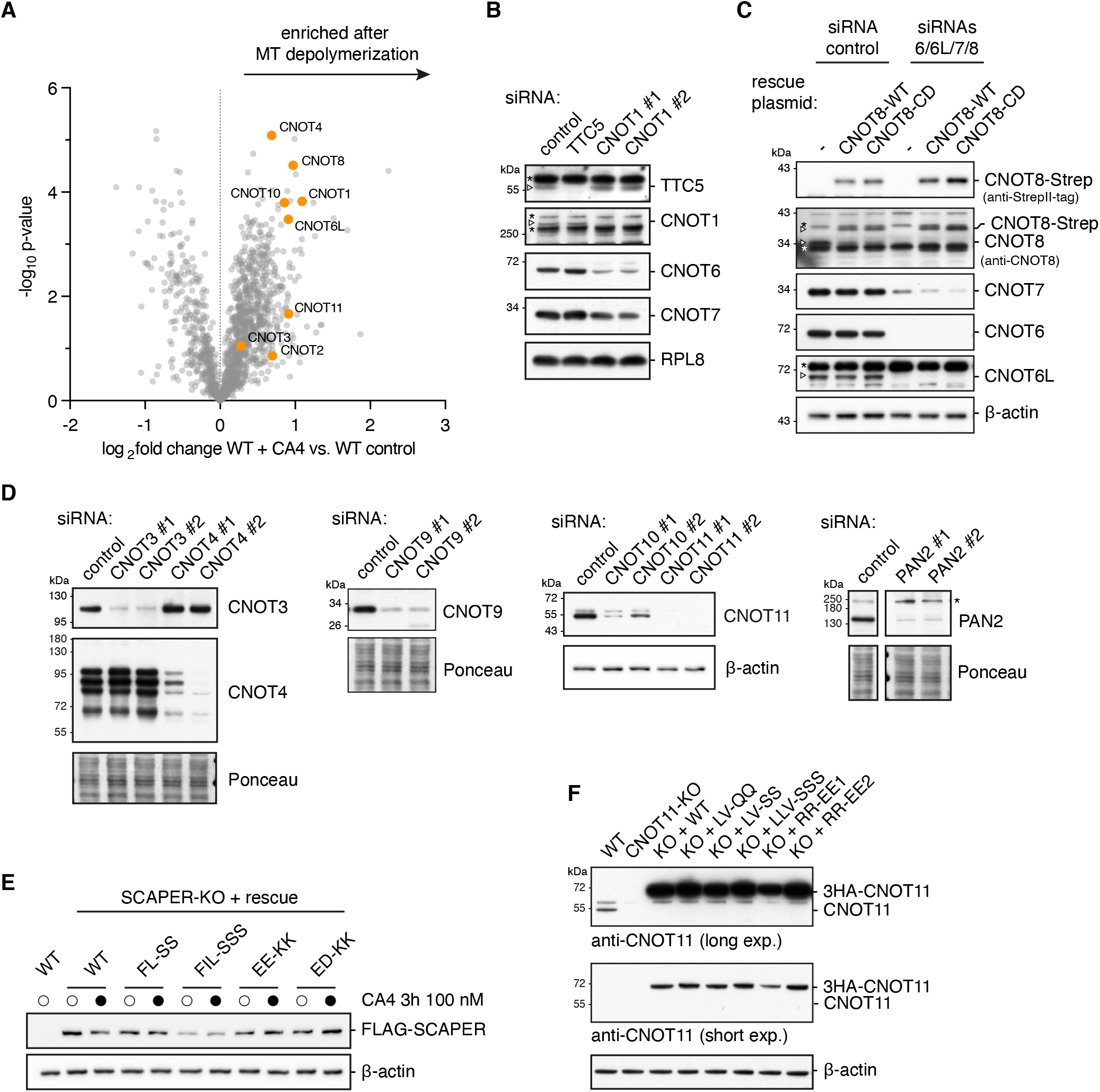
The CCR4-NOT complex is required for tubulin autoregulation, related to Fig. 4 and 5. (A) Proximity labelling using TurboID-SCAPER WT was performed in control conditions or after microtubule (MT) depolymerization with 100 nM Combretastatin A4 (CA4). Biotinylated proteins were enriched and analyzed by quantitative mass spectrometry. See also Supplementary Table S3. (B) Western blot analysis of samples as used in Fig. 4D confirming knockdown of CNOT1 and TTC5. Note that depletion of the CNOT1 scaffolding subunit also leads to destabilization of other CCR4-NOT subunits (CNOT6, CNOT7). Specific bands are indicated by triangles for TTC5 and CNOT1; non-specific bands are indicated by asterisks. (C) Western blot analysis of samples used in Fig. 4E. Signals from blots with an antibody against endogenous CNOT8 (second panel) show both CNOT8 depletion after knockdown and expression of siRNA-resistant rescue constructs CNOT8-WT and CNOT8-CD. Specific bands are indicated by triangles for CNOT8 and CNOT6L; non-specific bands are indicated by asterisks. (D) Western blot analysis of knockdown efficiency for selected samples used in Fig. 5A. Although we did not have access to a functional CNOT10 antibody, we note that CNOT10-KD leads to destabilization of CNOT11, corroborating that they might act as a functional unit. (E) Western blot analysis of expression levels of FLAG-SCAPER constructs used in Fig. 5D. FL-SS: F628S, L632S; FIL-SSS: F628S, I629K, L632S; EE-KK: E618K, E625K; ED-KK: E633K, D640K. (F) Western blot analysis of expression levels of CNOT11 and rescue constructs used in Fig. 5E. Membranes were probed with an antibody raised against a peptide of endogenous CNOT11. The epitope was not affected by the indicated mutations. LV-QQ: L405Q, V454Q; LV-SS: L405S, V454S; LLV-SSS: L405S, L451S, V454S; RR-EE1: R447E, R450E; RR-EE2: R461E, R485E.

**Figure S8.**
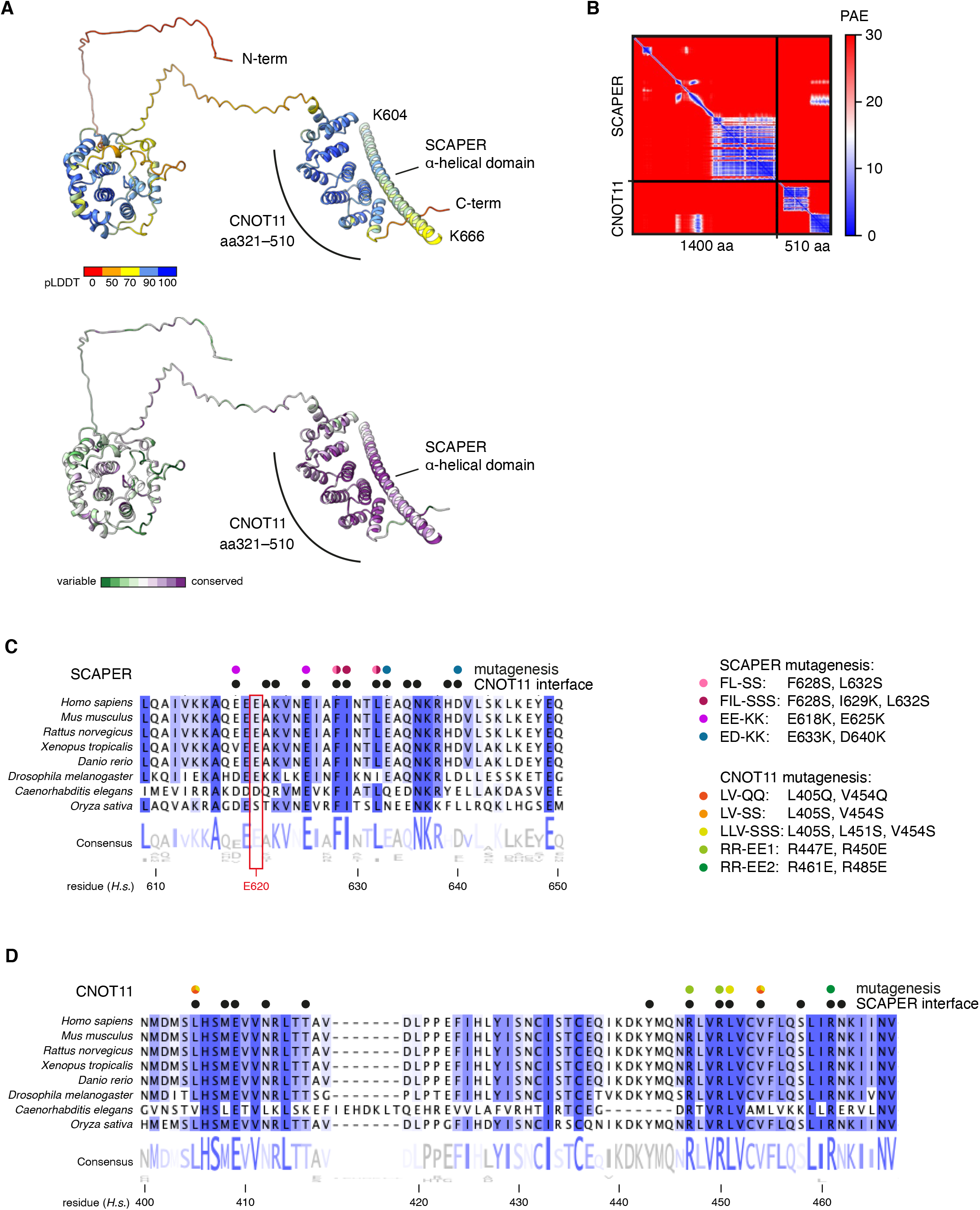
Structural prediction of a putative SCAPER-CNOT11 complex, related to Fig. 5. (A) Overview of an AF2 multimer predicted complex between CNOT11 and SCAPER coloured by predicted local distance difference test score (pLDDT, top, dark blue indicates high confidence), or per residue conservation (bottom). For SCAPER, only residues 604–666 of the α-helical domain are displayed for clarity. (B) PAE matrix of the model displayed in panel A. (C) Multiple sequence alignment of SCAPER homologs across the indicated species generated using Clustal Omega and displayed using Jalview software^84,85^. Highly conserved residues are shaded in blue. Residues in close proximity to CNOT11 in the AF2 predicted model are marked with black dots and residues targeted in mutagenesis experiments are marked with color-coded dots. The E620 residue is highlighted by a red box. (D) Multiple sequence alignment of CNOT11 homologs generated as in panel C. Residues in close proximity to SCAPER in the AF2 model are marked by black dots and residues targeted by mutagenesis are shown with coloured dots. Residue R485 is not shown in this alignment.

**Figure S9.**
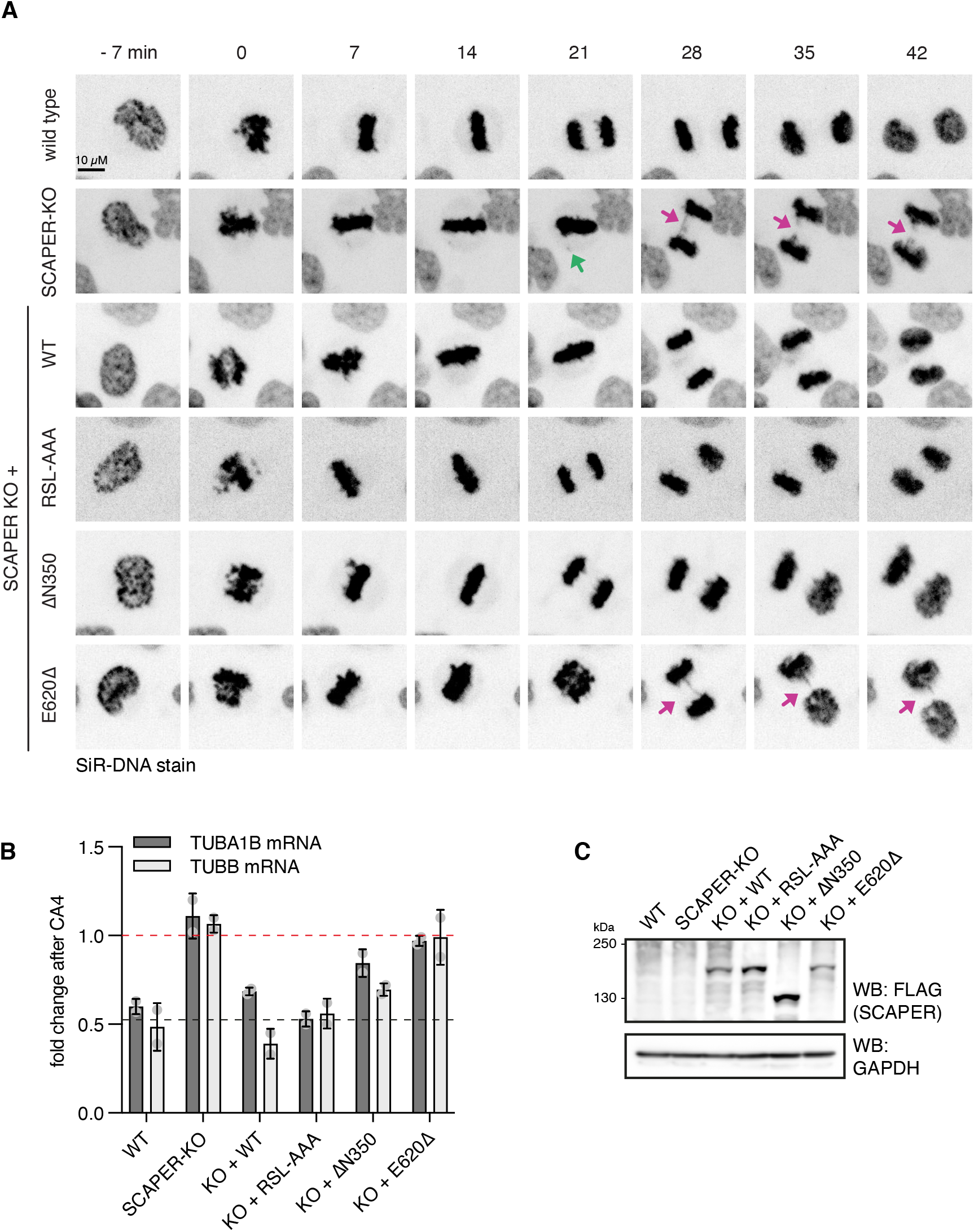
SCAPER is required for faithful mitosis, related to Fig. 6. (A) Examples from time-lapse imaging of HeLa cells of the indicated genotype going through mitosis. DNA was visualized using SiR-DNA stain and maximum intensity projections are shown. Frames were aligned to nuclear envelope breakdown (t = 0). Misaligned chromosomes and segregation errors are highlighted by green and magenta arrows, respectively. (B) Autoregulation assay for HeLa cell lines used in panel A and Fig. 6. (C) Total protein analysis by western blotting for FLAG-tagged SCAPER in the cell lines used in panel A and Fig. 6.

**Supplementary Table S1.** Cryo-EM data collection, processing, refinement and model statistics.

| **Data Collection** | Dataset 1 |
| --- | --- |
| Voltage (keV) | 300 |
| Pixel size (Å) | 0.829 |
| Detector | Gatan K3 |
| Defocus range (µm) | -1.2 to -2.7 |
| Electron dose (e<sup>-</sup> frame<sup>-1</sup> Å<sup>-2</sup>) | 1 |
| **Data Processing** |  |
| Useable micrographs | 19176 of 20932 |
| Particles picked | 1227269 |
| Final particles | 18949 |
| Map sharpening B-factor (Å<sup>2</sup>) | -20 |
| Resolution (Å) | 2.8 |
| EMDB accession code | EMD-16155 |
| PDB accession code | 8BPO |
| **Model Composition** |  |
| Chains | 62 |
| Non-hydrogen atoms | 143326 |
| Protein residues | 7633 |
| RNA bases | 3814 |
| Metals (Mg<sup>2+</sup>/Zn<sup>2+</sup>) | 214/5 |
| **Refinement** |  |
| Resolution (Å) | 2.9 |
| CC (mask) | 0.85 |
| **R.M.S deviations** |  |
| Bond lengths (Å) | 0.004 |
| Bond angles (°) | 0.851 |
| **Validation** |  |
| Molprobity score | 1.40 |
| Clashscore, all atoms | 3.79 |
| Rotamers outliers (%) | 0.03 |
| Cβ outliers (%) | 0.03 |
| **Ramachandran plot** |  |
| Favored (%) | 96.47 |
| Allowed (%) | 3.50 |
| Outliers (%) | 0.03 |

